# mTORC1 and JUN are activated after deletion of Prohibitin 1 in Schwann cells and may link mitochondrial dysfunction to demyelination

**DOI:** 10.1101/2020.11.25.398032

**Authors:** Gustavo Della-Flora Nunes, Emma R. Wilson, Edward Hurley, Bin He, Bert W. O’Malley, Yannick Poitelon, Lawrence Wrabetz, M. Laura Feltri

## Abstract

Schwann cell (SC) mitochondria are quickly emerging as an important regulator of myelin maintenance in the peripheral nervous system (PNS). However, the mechanisms underlying demyelination in the context of mitochondrial dysfunction in the PNS are incompletely understood. We recently showed that conditional ablation of the mitochondrial protein Prohibitin 1 (*Phb1*) in SCs causes a severe and fast progressing demyelinating peripheral neuropathy, but the mechanism that causes failure of myelin maintenance remained unknown. Here, we report that mTORC1 and JUN are continuously activated in the absence of *Phb1*, likely due to mitochondrial damage. Moreover, we demonstrate that these pathways are involved in the demyelination process, and that inhibition of mTORC1 using rapamycin partially rescues the demyelinating pathology. Therefore, we propose that mTORC1 and JUN may play a critical role as executioners of demyelination in the context of perturbations to SC mitochondria.

## 1. Introduction

Schwann cells (SCs) are the main glial cell type of the peripheral nerves, where they closely associate with axons (for review, see (Wilson et al., 2020)). Many axons extend very far from neuronal cell bodies, preventing fast delivery of essential cellular substrates. Therefore, SCs are believed to provide essential trophic and metabolic support to nearby axons (Nave, 2010). In addition, SCs identify axons larger than 1μm in diameter and wrap them in multiple layers of a specialized membrane extension known as myelin. The myelin sheath reduces the capacitance of the axonal membrane, and its discontinuous structure enables “saltatory conduction”, whereby ionic exchanges are concentrated in small myelin-free regions called nodes of Ranvier. The importance of SCs is evident from the great variety of inherited and acquired peripheral neuropathies that arises when these cells are impaired (England and Asbury, 2004).

Myelin is intuitively perceived as a stable structure, which can be exemplified by the remarkable discovery of preserved myelin ultrastructure in a 5000-year-old ice man (Hess et al., 1998). This notion of stability was initially confirmed by studies investigating the turnover of myelin components in brain, with many myelin proteins and lipids showing half-lives of months (Smith and Eng, 1965, Fischer and Morell, 1974). Nevertheless, even early studies were quick to point out that some of the myelin components had much faster turnover rates (Singh and Jungalwala, 1979, Sabri et al., 1974, Hayes and Jungalwala, 1976), suggesting that portions of the myelin (especially its non-compact regions) could be more dynamic. Furthermore, we now know that maintenance of the myelin structure is not passive, requiring sustained expression of the transcription factors EGR2 (also known as Krox20) (Decker et al., 2006) and SOX10 (Bremer et al., 2011) in SCs, as well as continuous synthesis of myelin proteins (Meschkat et al., 2020) and lipids (Zhou et al., 2020) in brain.

Recently, mitochondria and cell metabolism were also implicated in myelin formation and maintenance in SCs. We reported that ablation of the primarily mitochondrial protein prohibitin 1 (*Phb1*) in SCs greatly impairs myelin maintenance in the PNS (Della-Flora Nunes et al., 2020). In addition, SC-specific deletion of the mitochondrial transcription factor *Tfam* (Viader et al., 2011), the respiratory chain component *Cox10* (Funfschilling et al., 2012), the metabolic regulator *Lkb1* (Beirowski et al., 2014, Pooya et al., 2014, Shen et al., 2014), the NAD+ synthetizing enzyme *Nampt* (Sasaki et al., 2018), or the nutrient-sensing O-linked N-acetylglucosamine transferase *Ogt* (Kim et al., 2016), all lead to peripheral neuropathy phenotypes in mice. Nonetheless, the mechanisms linking mitochondrial dysfunction to impaired myelin maintenance in SCs remain poorly understood.

Here we report that mice lacking *Phb1* in SCs (Phb1-SCKO) activate a response involving JUN and the mTORC1 pathway, possibly directly downstream of the mitochondrial damage. In addition, both JUN and mTORC1 seem to participate in the demyelination process in Phb1-SCKO animals. Moreover, inhibition of mTORC1 using rapamycin is able to partially rescue morphological and functional aspects of the phenotype of Phb1-SCKO mice. These results reveal a previously unknown mechanism that contributes to myelin loss secondary to SC mitochondrial damage. Furthermore, our findings implicate JUN and mTORC1 in the SC adaptation to mitochondrial dysfunction, raising the possibility that JUN and mTORC1 may also be involved in the response to mitochondrial damage in other cell types.

## 2. Results

### 2.1. Deletion of *Phb1* in SCs elicits fast and widespread myelin loss in the PNS

We recently reported that deletion of *Phb1* specifically in SCs causes a severe and fast progressing demyelinating phenotype in mice (Della-Flora Nunes et al., 2020). Mice lacking *Phb1* in SCs (Phb1-SCKO) show the first signs of myelin loss at postnatal day 20 (P20), with demyelination peaking around P40-P60 and causing partial or complete hindlimb paralysis around P90 (Della-Flora Nunes et al., 2020) (**Figures 1A** **and** **1B**). Concomitant with the demyelination, Phb1-SCKO mice also display axonal degeneration, which follows a similar temporal progression (Della-Flora Nunes et al., 2020). This pathology is likely caused by the profuse changes in mitochondrial morphology and function that follow deletion of *Phb1* in SCs (Della-Flora Nunes et al., 2020). Nevertheless, the molecular mechanism linking mitochondrial damage to myelin destruction in Phb1-SCKO mice remains unknown.

**Figure 1.**
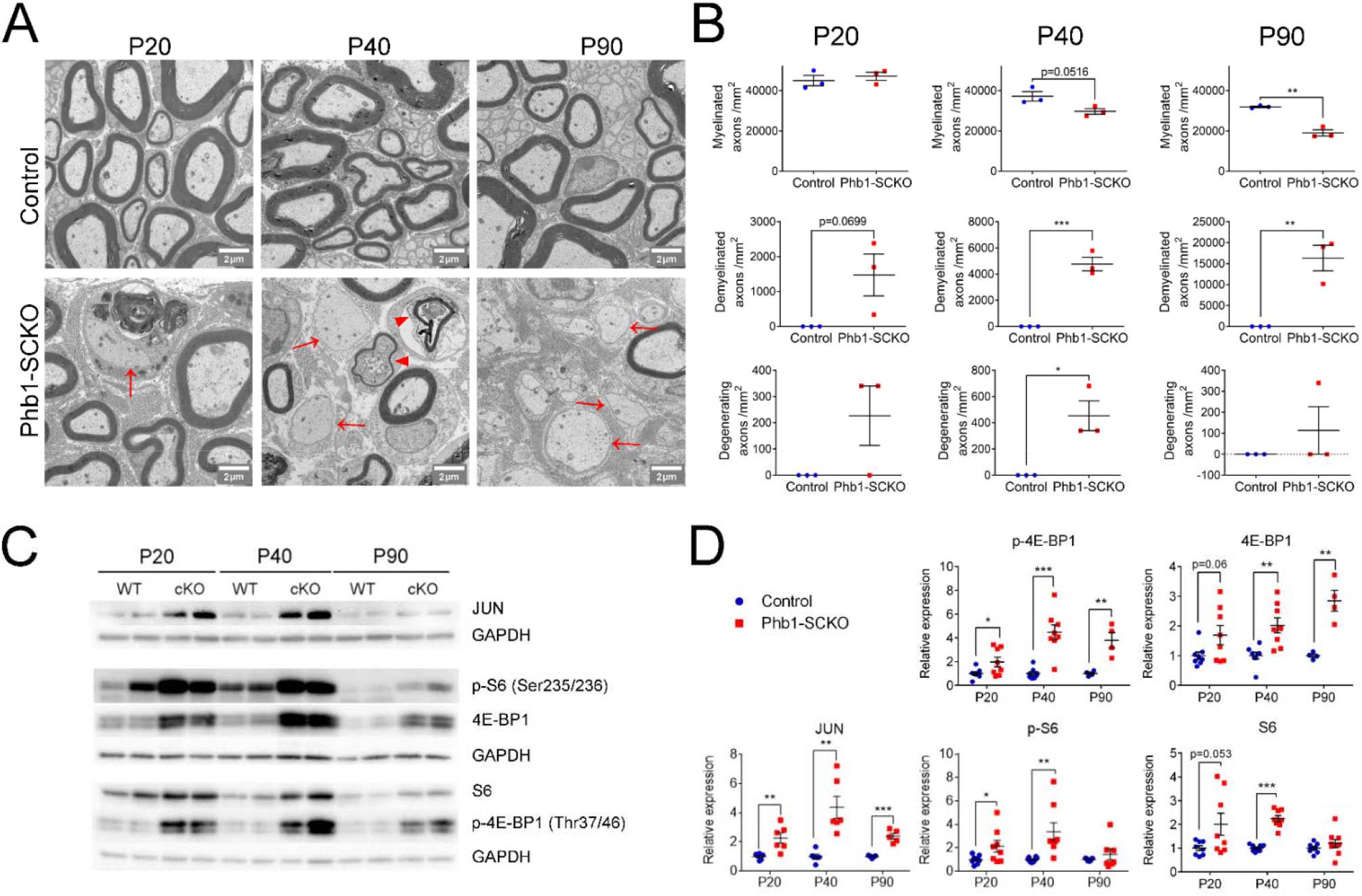
The mTORC1 and JUN pathways are activated upon deletion of Phb1. (**a**) Representative electron micrographs of sciatic nerves demonstrating the demyelinating phenotype of Phb1-SCKO mice. Note the presence of demyelinated axons (arrows) and degenerating axons (arrowheads). (**b**) Quantification of myelinated (top), demyelinated (center) and degenerating (bottom) axons at the analyzed ages. There is progressive demyelination and axonal degeneration in the sciatic nerves of Phb1-SCKO animals. N=3 animals per genotype. Unpaired two-tailed t-test. Myelinated axons [P20 (t=0.6619, df=4), P40 (t=2.745, df=4), P90 (t=8.164, df=4); demyelinated axons [P20 (t=2.457, df=4), P40 (t=9.165, df=4), P90 (t=5.322, df=4); degenerating axons [P20 (t=2.000, df=4), P40 (t=4.000, df=4), P90 (t=1.000, df=4) (**c**) Representative western blot from sciatic nerve lysates reveal that deletion of *Phb1* leads to upregulation of JUN and increased phosphorylation of the mTORC1 targets S6 and 4E-BP1. (**d**) Quantitative analysis of the relative expression of the proteins in (c). N=4-9 animals per genotype. Unpaired two-tailed t-test. p-4E-BP1 [P20 (t=2.264, df=14), P40 (t=5.337, df=14), P90 (t=4.228, df=6)]; 4E-BP1 [P20 (t=2.013, df=14), P40 (t=3.625, df=14), P90 (t=5.199, df=6)]; JUN [P20 (t=3.53, df=10), P40 (t=4.34, df=10), P90 (t=7.172, df=8)]; p-S6 [P20 (t=2.186, df=14), P40 (t=3.086, df=14), P90 (t=0.936, df=12)]; S6 [P20 (t=2.104, df=14), P40 (t=8.838, df=14), P90 (t=1.084, df=16)]; * p<0.05, ** p<0.01, *** p<0.001.

### 2.2. Ablation of *Phb1* in SCs leads to upregulation of JUN and activation of the mTORC1 pathway

Although little is known about how myelin is maintained long-term, it seems that preservation of myelin with the correct structure and thickness is an active process, requiring the continuous activation of certain cellular machinery in SCs (Bremer et al., 2010, Zhou et al., 2020, Meschkat et al., 2020, Bremer et al., 2011, Decker et al., 2006). We therefore hypothesized that the SC response to mitochondrial dysfunction may inadvertently interfere with pathways critical for myelin maintenance. Seeking to find the molecular link between compromised mitochondrial function and demyelination, we investigated the status of the following molecular pathways previously reported to play roles in preservation of myelin: mTORC1 (important for myelination (Beirowski et al., 2017, Figlia et al., 2017) and remyelination (Norrmen et al., 2018)), ERK 1/2 (whose activation in sufficient to trigger demyelination (Napoli et al., 2012)), AKT (important to regulate myelin sheath thickness (Domenech-Estevez et al., 2016)), JUN (the master transcription factor of nerve repair (Arthur-Farraj et al., 2012)), and eIf2α (the core protein in the integrated stress response, ISR, which is important for myelin maintenance in the context of perturbed protein homeostasis in SCs (Scapin et al., 2020, D’Antonio et al., 2013)).

We had previously reported that phosphorylation of eIf2α is protective in Phb1-SCKO mice, and that they only show minor changes in the ERK 1/2 pathway, which are unlikely to be sufficient to initiate demyelination (Della-Flora Nunes et al., 2020). The same is true for the AKT pathway (**Supplementary Figure 1**). On the other hand, Phb1-SCKO animals show continuous upregulation of JUN and activation of the mTORC1 pathway, as measured by the protein and phosphorylation levels of the downstream targets 4E-BP1 and S6 (**Figures 1C** **and** **1D**). Importantly, these changes are evident even before overt demyelination.

Both JUN and mTORC1 are required for formation of repair SCs after nerve injury, a cell type that mounts a response that favors axon regrowth, tissue reinnervation and clearance of myelin and other debris (Jessen and Mirsky, 2019). In the case of myelinating SCs, this transdifferentiation to the repair phenotype involves myelin removal followed by its autophagic degradation (myelinophagy) (Gomez-Sanchez et al., 2015). This, associated to the fact that enforced JUN expression is sufficient to trigger demyelination (Fazal et al., 2017), raises the possibility that JUN and mTORC1 may underly important aspects of the peripheral neuropathy that ensues in Phb1-SCKO mice.

### 2.3. mTORC1 and JUN can be activated downstream of mitochondrial perturbations in SCs

Both JUN and mTORC1 have been previously implicated in the cellular response to mitochondrial dysfunction. The mitochondrial unfolded protein response (mtUPR) is believed to be partly regulated by binding of an AP-1 transcription factor (postulated to be JUN) to the promoter of CHOP and C/EBPβ (Horibe and Hoogenraad, 2007). On the other hand, mTORC1 was shown to be upstream of the integrated stress response (ISR) in muscle, regulating the progression of a mitochondrial myopathy (Khan et al., 2017).

For this reason, we sought to investigate if JUN and mTORC1 were activated downstream of perturbations to SC mitochondria. We treated primary rat SCs with compounds that affect different aspects of mitochondrial function: Carbonyl cyanide-p-trifluoromethoxyphenylhydrazone (FCCP), an ionophore that dissipates the mitochondrial membrane potential; Oligomycin, an inhibitor of the ATP synthase; or Antimycin A, an inhibitor of mitochondrial complex III. We then evaluated the expression levels of JUN and downstream targets of the mTORC1 pathway (4E-BP1 and S6), as well as components of the other pathways we previously demonstrated to be altered in Phb1-SCKO mice (Della-Flora Nunes et al., 2020): eIF2α (which is phosphorylated downstream of different cellular stressors to activate the ISR), BiP (a chaperone also known as HSPA that is upregulated in response to stress in the endoplasmic reticulum, ER), and Opa1 (a protein essential for mitochondrial fusion that, under situations of mitochondrial damage, is cleaved proteolytically, inhibiting mitochondrial fusion). Short-time treatment (24h) with these compounds resulted only in minor changes in these pathways (**Supplementary Figures 2A and 2B**). Cells treated with FCCP for 24h presented with elevated levels of p-eIF2α and reduced ratio between the long and short isoforms of Opa1 (**Supplementary Figures 2A and 2B**). This goes in line with previous reports showing that the ISR can quickly be activated directly downstream of compromised mitochondria (Viader et al., 2013, Mick et al., 2020). Oligomycin and Antimycin A applied to SCs for 24h were unable to elicit significant changes in the conditions tested. On the other hand, a seven-day treatment of SCs with Oligomycin or Antimycin A triggered robust stimulation of the ISR (**Figures 2A** **and** **2B**). Furthermore, this lengthier treatment regimen also induced a potent activation of the mTORC1 pathway (**Figures 2A** **and** **2B**). Hence, it is possible that the mTORC1 pathway participates in the SC adaptation to long-term mitochondrial impairments.

**Figure 2.**
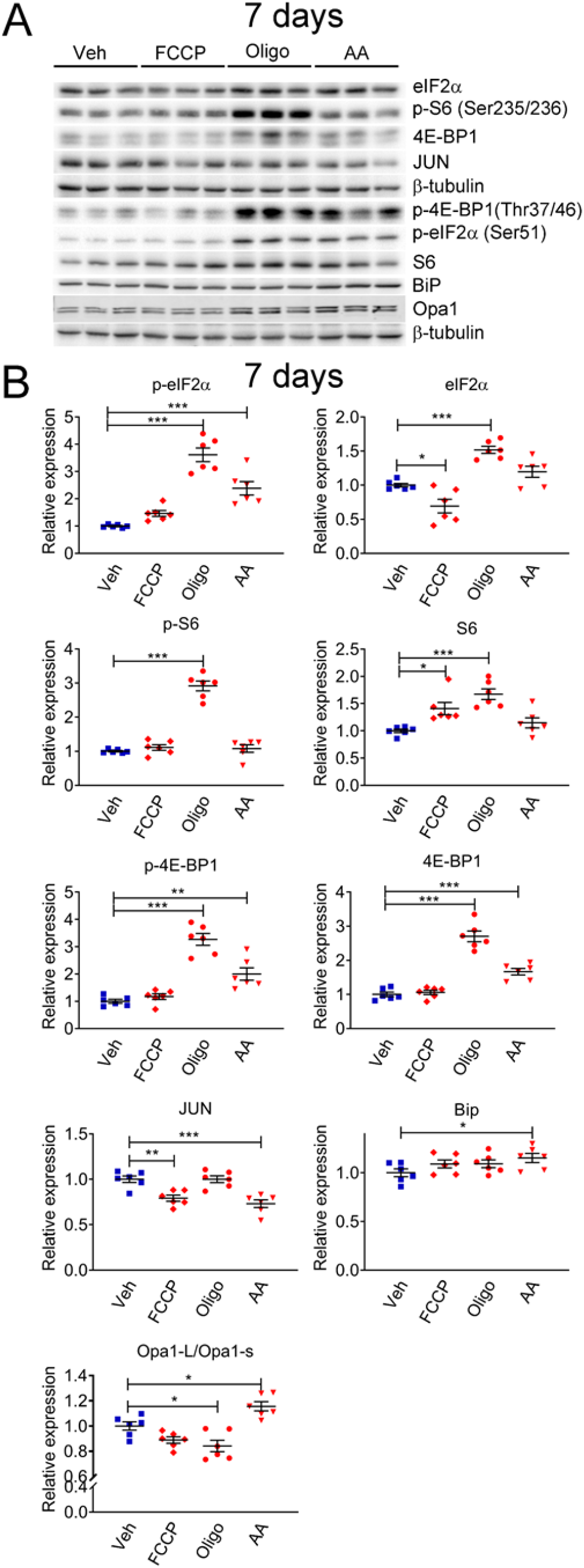
Mitochondrial perturbations in SCs *in vitro* lead to activation of the mTORC1 pathway. (**a**) Representative western blot from cell lysates of primary rat SCs treated with 5 μM FCCP, 2.5 μM oligomycin (Oligo), 10 μM antimycin A (AA) or vehicle (Veh) for seven days. There is a robust activation of the ISR when SCs are treated long-term with oligomycin or antimycin A, conditions that also result in activation of th mTORC1 pathway. (**b**) Quantification of th experiments in (c). N=6 wells per condition. One-way ANOVA corrected for multipl comparisons with the Dunnett method. F (3, 20) p-eIF2α = 39.18, p<0.0001; F (3, 20) eIF2α = 23.9, p<0.0001; F (3, 20) p-S6 = 87.39, p<0.0001; F (3, 20) S6 = 11.13, p<0.0001; F (3, 20) p-4E-BP1 = 37.53, p<0.0001; F (3, 20) 4E-BP1 = 59.55, p<0.0001; F (3, 20) JUN = 13.82, p<0.0001; F (3, 20) Bip = 2.211, p=0.118; F (3, 20) Opa1-L/Opa1-s = 15.09, p<0.0001. * p<0.05, ** p<0.01, *** p<0.001.

Next, we interrogated whether JUN or mTORC1 activity were associated with presence of compromised mitochondrial network *in vivo*. For this experiment, we analyzed Phb1-SCKO animals in which SC mitochondria were genetically labelled with the PhAM reporter. The PhAM mouse line contains a floxed STOP construct coding for a mitochondrially-targeted version of the Dendra2 fluorophore (Pham et al., 2012). We previously reported that, at P40, about 20% of SCs in Phb1-SCKO mice show disrupted mitochondrial network, especially away from the cell body, where PhAM is almost completely undetectable (Della-Flora Nunes et al., 2020). Staining of individual myelinated fibers with JUN indicated that JUN immunoreactivity was significantly associated with the presence of compromised mitochondria (**Figures 3A** **and** **3B**). Moreover, 78.81% of all SCs with damaged mitochondria showed strong JUN nuclear expression, while only 23.15% of SCs with reasonably well-preserved mitochondria stained positive for JUN (**Figure 3C**). In contrast, we did not identify any association between mitochondrial loss and expression of p-S6 (**Figures 3D-F**).

**Figure 3.**
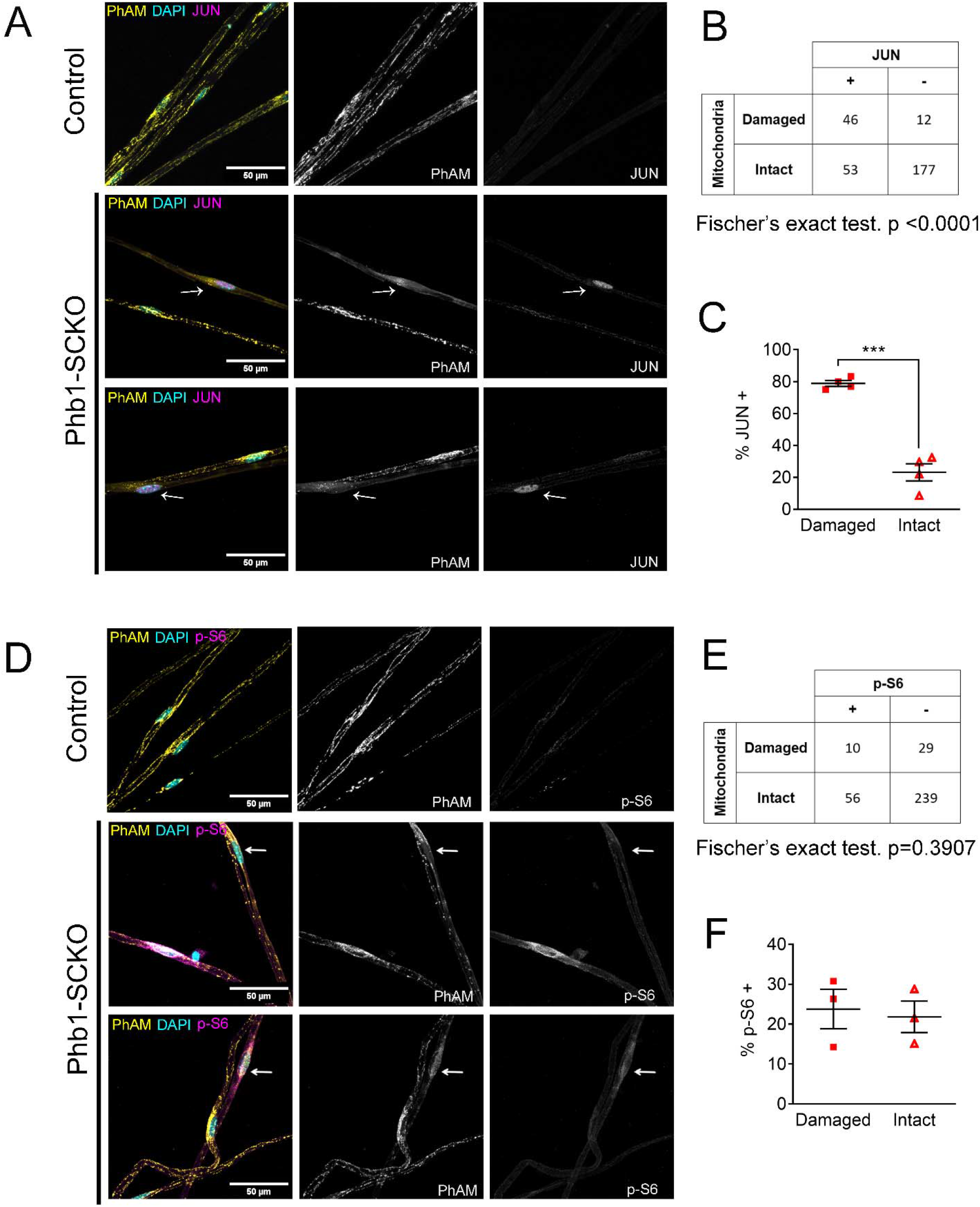
JUN expression is associated with mitochondrial loss *in vivo*, but mTORC1 activation is not. (**a**) Immunofluorescence for JUN in teased fibers from sciatic nerves of P40 animals expressing PhAM in SC mitochondria. Note that SCs of Phb1-SCKO mice that show perturbation to their mitochondrial network tend to also show high nuclear JUN expression (magenta). (**b**) There is an association between mitochondrial damage and JUN staining in SCs of Phb1-SCKO mice. N= 4 animals. Fischer’s exact test. (**c**) The majority of SCs of Phb1-SCKO mice that lack PhAM expression away from the cell body show positive staining for JUN. N= 4 animals. Paired two-tailed t-test (t=14.09, df=3) (**d**) Immunofluorescence for phosphorylated S6 ribosomal protein (p-S6) in teased fibers from sciatic nerves of animals expressing PhAM. Arrows show two SCs with damaged mitochondria, the one on top was considered p-S6 −, while the one on the bottom was classified as p-S6 +. (**e**) There is no correlation between mitochondrial damage and p-S6 staining in SCs of Phb1-SCKO mice. N= 3 animals. Fischer’s exact test. (**f**) The percentage of SCs labelled with p-S6 is constant regardless of the status of their mitochondrial network visualized by PhAM. N= 4 animals. Paired two-tailed t-test (t=0.236, df=2). *** p<0.001.

Thus, our data supports the hypothesis that both JUN and mTORC1 can be activated downstream of perturbations to mitochondria under certain circumstances, raising the possibility that JUN and mTORC1 may participate in the response to mitochondrial stress in SCs. The discrepancy in our *in vitro* and *in vivo* data may be because of the already elevated JUN expression in SCs *in vitro* (SCs in culture show an immature phenotype (Schmid et al., 2014, Morgan et al., 1991)). In addition, mitochondrial loss is a late event *in vivo*, while mTORC1 activation occurs relatively quickly, after only one week *in vitro*. Thus, mTORC1 and JUN may be activated in different stages of the SC response to mitochondrial damage, with mTORC1 preceding JUN temporally.

### 2.4. JUN and mTORC1 are associated with demyelination in Phb1-SCKO mice

Given the possibility that JUN and mTORC1 could be activated downstream of mitochondrial dysfunction, and the importance of these pathways for formation of repair SCs, we asked if demyelination in Phb1-SCKO mice was preferentially happening in SCs with overactive JUN or mTORC1. With this goal, we immunostained individual myelinated fibers of Phb1-SCKO animals for myelin proteins (P0 and MBP) in conjunction with JUN or p-S6. We then identified SCs undergoing demyelination by the presence of myelin fragments inside SCs (myelin ovoids). In our analysis, all of the evaluated fibers containing myelin ovoids showed intense nuclear JUN staining (**Figures 4A-C**), suggesting a strong association between JUN expression and demyelination. Similarly, we also identified a correlation between p-S6 and the presence of myelin ovoids (**Figures 4D** **and** **4E**), and a greater proportion of the SCs containing myelin ovoids were also positive for p-S6, although this difference did not reach statistical significance (**Figure 4F**).

**Figure 4.**
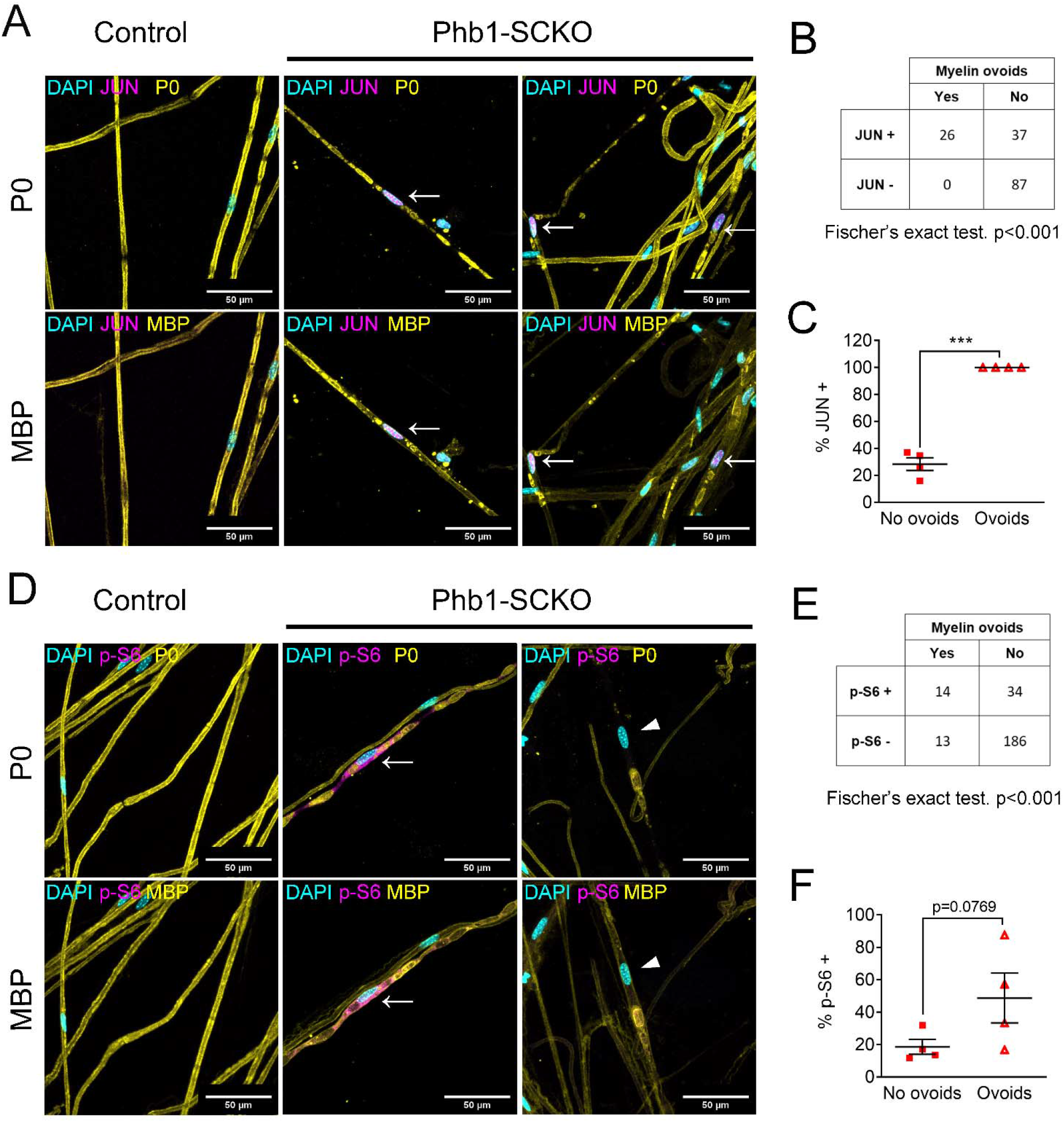
Activation of the mTORC1/JUN axis is associated with demyelination. (**a**) Teased fibers from tibial nerves of 40-day-old mice stained for myelin proteins (MBP and P0) and JUN. All cells containing myelin ovoids were also JUN positive (arrows). (**b**) JUN immunoreactivity and presence of myelin ovoids are associated. N= 4 animals. Fischer’s exact test. (**c**) Cells that present with myelin ovoids show a higher percentage of JUN immunoreactivity. N= 4 animals. Paired two-tailed t-test (t=15.05, df=3). (**d**) Teased fibers from tibial nerves of 40-day-old mice stained for myelin proteins (MBP and P0) and phosphorylation of S6, a downstream target of mTORC1. Arrows and arrowheads show cells containing myelin avoids and that are p-S6 positive and p-S6 negative, respectively. (**e**) There is an association between p-S6 reactivity and presence of myelin ovoids. N= 4 animals. Fischer’s exact test. (**f**) The percentage of cells positive for p-S6 tends to be higher among cells that present with myelin ovoids. N= 4 animals. Paired two-tailed t-test (t=2.651, df=3). *** p<0.001.

We then tested if Phb1-SCKO mice showed activation of other pathways known to participate in breakdown and degradation of myelin. First, we probed Phb1-SCKO animals for molecules downstream of JUN and found the mRNA expression of multiple JUN targets to be altered in the expected direction: upregulation of the neurotrophin Glial-derived neurotrophic factor (*Gdnf*) and the signaling molecule Sonic hedgehog (*Shh*); downregulation of the adhesion molecule Cadherin-1 (*Cdh1*) and of the myelin genes Myelin basic protein (*Mbp*) and Myelin protein zero (*Mpz*, which codes for the P0 protein) (**Supplementary Figure 3A**). Phb1-SCKO mice also showed an overexpression of Mixed lineage kinase domain-like (MLKL), a protein recently implicated in dismantling the myelin sheath after nerve injury (Ying et al., 2018) (**Supplementary Figures 3B and 3D**). Moreover, deletion of *Phb1* in SCs also resulted in upregulation of the autophagic machinery, which is important for myelinophagy (**Supplementary Figures 3C and 3D**).

Taken together, these results indicate that Phb1-SCKO mice show an overactive myelin breakdown machinery, which includes mTORC1 and JUN, whose activation is associated with demyelination. Given these results, it is tempting to hypothesize that JUN and mTORC1 can be key pathways orchestrating the demyelination process in Phb1-SCKO mice.

### 2.5. The ISR has minor effects on the other evaluated pathways

We recently showed that the ISR is likely a beneficial response in Phb1-SCKO mice (Della-Flora Nunes et al., 2020). Activation of the ISR frequently leads to alterations in the mTORC1 pathway (Ryoo and Vasudevan, 2017, Zhang et al., 2019). Thus, we asked whether the ISR is upstream of the pathways analyzed in this study. Inhibition of the ISR using a daily injection of 2.5 mg/Kg ISRIB as previously reported (Della-Flora Nunes et al., 2020) did not significantly alter p-S6, JUN or Opa1 expression in Phb1-SCKO mice (**Supplementary Figure 4**). Nonetheless, this treatment was sufficient to result in a small reduction in levels of p-4E-BP1 and 4E-BP1 (**Supplementary Figures 4B and 4C**).

### 2.6. JUN may participate in the demyelination process, but ablation of JUN in Phb1-SCKO mice is not sufficient to ameliorate the neuropathy phenotype

In order to test the hypothesis that JUN coordinates demyelination in Phb1-SCKO mice, we crossed those animals to JUN floxed mice. At P40, presence of one (Phb1-SCKO; JUN Het) or two *Jun* floxed alleles (Phb1; JUN SCKO) resulted in a significant and dose-dependent reduction in JUN protein levels in sciatic nerves (**Figure 5A**). As expected, this reduction in JUN also led to a dose-dependent reduction in the number of demyelinated fibers and myelinophagy events (**Figures 5B** **and** **5C**). Nevertheless, deletion of *Jun* in Phb1-SCKO mice also caused a dose-dependent reduction in the number of myelinated axons in tibial nerves (**Figures 5B** **and** **5C**). In addition, nerves of Phb1; JUN SCKO contained bundles with large non-myelinated axons and a visible division of the nerve in smaller fascicles (hyper-fasciculation) (**Figures 5B-5D**). These phenotypes suggest that ablation of *Jun* in SCs already lacking *Phb1* may exacerbate the mild developmental defects that we previously observed in Phb1-SCKO mice (Della-Flora Nunes et al., 2020). As a consequence, *Jun* deletion is not sufficient to rescue the motor deficits of Phb1-SCKO mice in the rotarod (**Figure 5E**).

**Figure 5.**
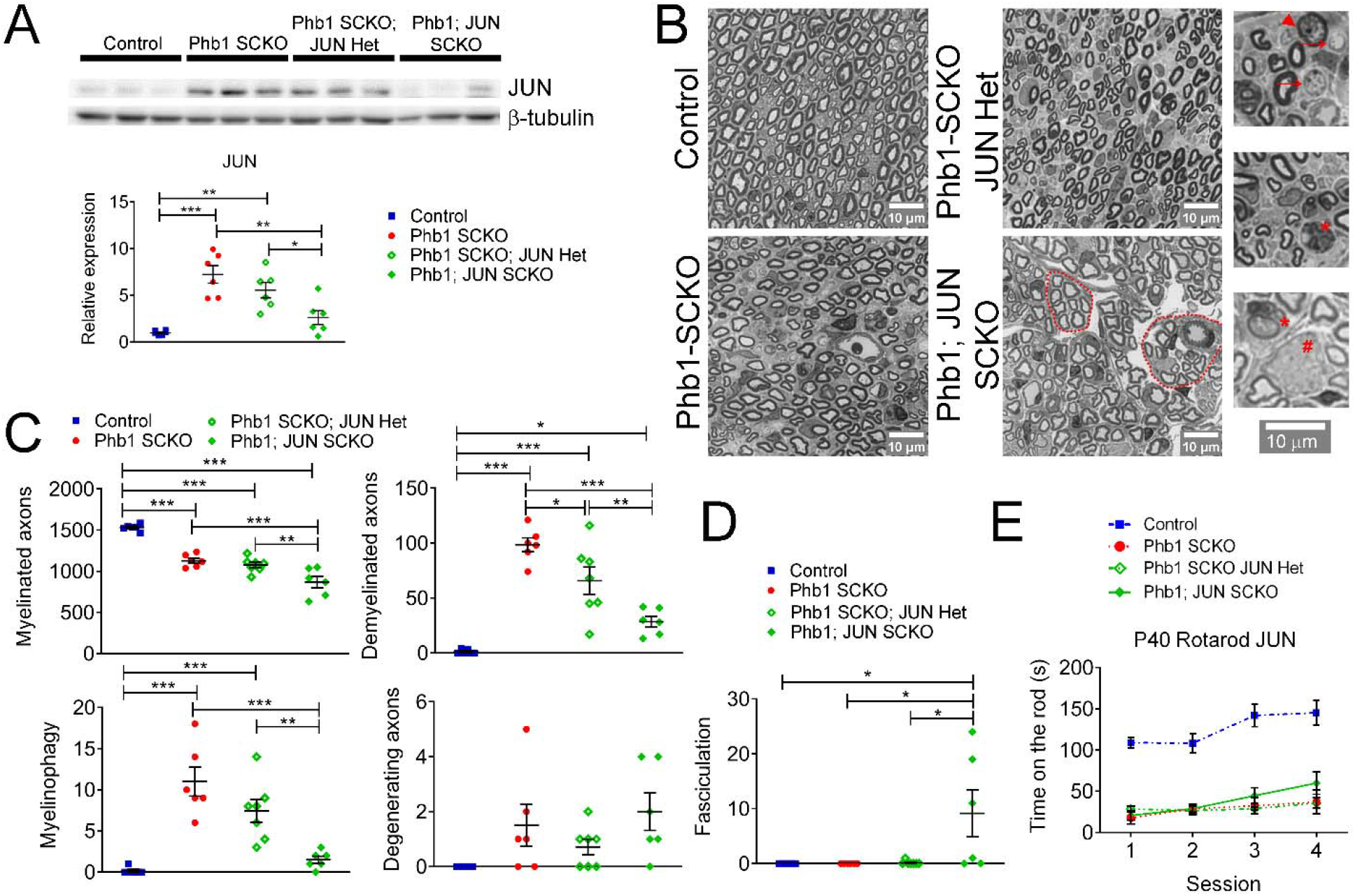
JUN participates in the demyelination process in Phb1-SCKO mice, but JUN ablation is unable to ameliorate the behavioral phenotype. (**a**) Top: Western blot from sciatic nerve lysates illustrating the reduction in JUN levels when SCs of P40 Phb1-SCKO mice have one (Phb1 SCKO; JUN Het) or both JUN alleles deleted (Phb1; JUN SCKO). Bottom: Quantification of the experiment represented in the top panel. N=6 animals per group. One-way ANOVA corrected for multiple comparisons using the Holm-Sidak method. F (3, 20) = 15.08, p<0.0001. (**b**) Representative semithin sections from tibial nerves. Note that nerves of Phb1; JUN SCKO mice show a division of axons into smaller fascicles (dotted lines), an abnormality known as hyper-fasciculation. Nerves of these animals also frequently have large bundle structures containing axons of mixed caliber, indicative of a mild radial sorting defect (see inset). Insets: Representative images of demyelinated axons (arrows), degenerating axon (arrowhead), myelinophagy (stars) and large bundles with axons of mixed caliber (pound). (**c**) Quantification of morphological parameters from semithin images reveal that JUN ablation results in a reduction in demyelinated axons and myelin degradation (myelinophagy; visualized by the presence of cytoplasmic myelin debris in SCs) in Phb1-SCKO mice. Nonetheless, animals in which JUN has been deleted show a dose-dependent reduction in myelinated axons, suggesting that deletion of JUN may amplify the developmental defect observed in Phb1-SCKO animals. N=6-7 animals per group. One-way ANOVA corrected for multiple comparisons using the Holm-Sidak method. F (3, 21) myelinated = 3.211, p=0.0438; F (3, 21) demyelinated = 5.064, p=0.0085; F (3, 21) myelinophagy = 2.667, p=0.074; F (3, 21) degenerating = 2.69, p=0.0724. (**d**) Phb1; JUN SCKO mice commonly show hyper-fasciculation. N=6-7 animals per group. One-way ANOVA corrected for multiple comparisons using the Holm-Sidak method. F (3, 21) = 16.67, p<0.001 (**e**) Deletion of JUN has no observable effect in the performance of Phb1-SCKO mice in the rotarod. N=6 animals per group. Two-way ANOVA corrected for multiple comparisons using the Holm-Sidak method. F (9, 60) interaction = 2.654, p=0.0117; F (3, 60) time = 19.65, p < 0.0001; F (3, 20) group = 33.63, p < 0.0001. * p<0.05, ** p<0.01, *** p<0.001.

To investigate if JUN was important to modulate other pathways of interest, we probed the protein levels of eIF2α, p-eIF2α, 4E-BP1, p-4E-BP1, S6, p-S6, Opa1 and BiP. Deletion of *Jun* in Phb1-SCKO mice had a dose-dependent effect on levels of p-4E-BP1 and p-S6, suggesting that JUN upregulation may be partially responsible for mTORC1 overactivation (**Supplementary Figures 5A and 5B**). A similar effect was observed on BiP levels (**Supplementary Figures 5A and 5B**). JUN, however, does not seem to be important to regulate the ISR, since p-eIF2α levels are unaltered in Phb1-SCKO mice upon *Jun* deletion (**Supplementary Figures 5A and 5B**). In agreement with this finding, ablation of *Jun* is also unable to alter the mRNA levels of ATF4 targets (*Asns*, *Chac1*, *Pck2* and *Ddit3*, also known as *Chop*), which are upregulated downstream of p-eIF2α during ISR in Phb1-SCKO mice (**Supplementary Figures 5C**).

Mice containing two floxed *Jun* alleles and one floxed *Phb1* allele (JUN SCKO; Phb1 Het) are statistically indistinguishable from control mice in all of the aforementioned analyses (**Supplementary Figure 6)**, indicating that deletion of JUN alone is unable to elicit any of the changes observed in Phb1-SCKO; JUN Het or Phb1; JUN SCKO mice.

### 2.7. Overactivation of mTORC1 is causal for demyelination in Phb1-SCKO mice

Next, we sought to explore the role of mTORC1 in the pathology observed in Phb1-SCKO mice. Considering the important developmental role of mTORC1 (Beirowski et al., 2017, Figlia et al., 2017), it would be necessary to inhibit mTORC1 after myelination is completed.

Therefore, we opted for a pharmacological treatment instead of a genetic approach. We administered Phb1-SCKO mice and controls with daily injections of the well-established mTORC1 inhibitor rapamycin from P20 to P40 (**Figure 6A**). Rapamycin binds to FK506-binding protein (FKBP12), which becomes an allosteric inhibitor of mTORC1 (Li et al., 2014) (**Figure 6A**). Rapamycin treatment was efficient, and significantly reduced the levels of p-4E-BP1 and p-S6 in sciatic nerves of P40 Phb1-SCKO mice (**Figure 6B** **and** **6C**). Interestingly, suppression of mTORC1 activity resulted in almost complete rescue of nerve morphology in Phb1-SCKO animals at P40. Mutant mice treated with rapamycin had the same number of myelinated axons, demyelinated axons, degenerating axons and myelinophagy as their littermate controls (**Figures 6D** **and** **6E**). Deletion of *Phb1* in Schwann cells did not alter myelin thickness on the myelinated axons that remained, and this was unaffected by rapamycin treatment (**Figure 6F**). Consistent with the results of our morphological analyses, rapamycin was also able to partially ameliorate the nerve conduction velocity in Phb1-SCKO mice (**Figure 6G**). Nonetheless, rapamycin could not improve the amplitude of the compound muscle action potential (CMAP) of mutant mice (**Figure 6G**) nor the motor deficits of Phb1-SCKO mice measured by the rotarod (**Figure 6H**). In summary, these results suggest that mTORC1 overactivation is key to induce demyelination in Phb1-SCKO mice, and that inhibition of the mTORC1 pathway can provide meaningful benefit to nerve conduction velocity.

**Figure 6.**
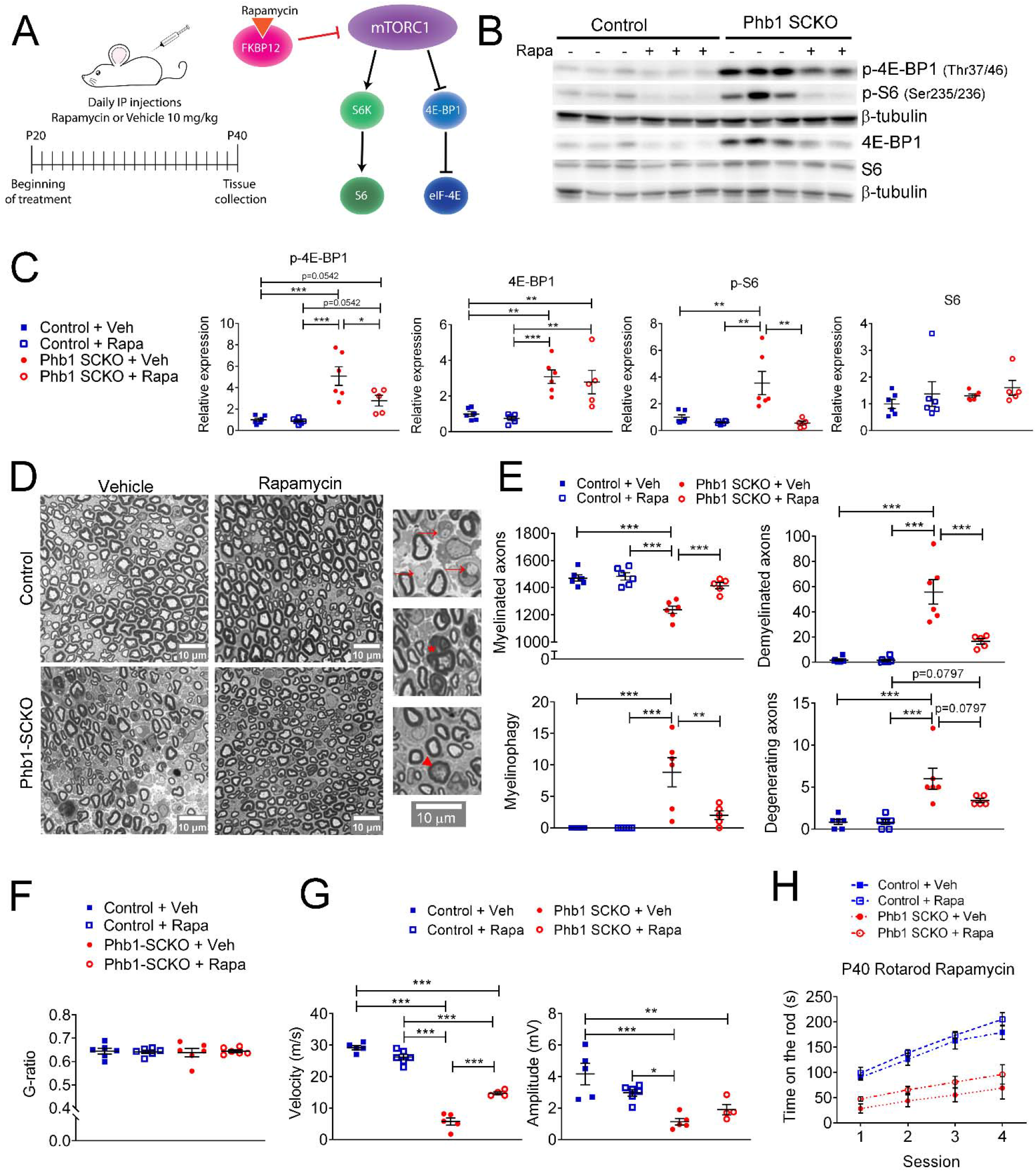
Inhibition of mTORC1 prevents demyelination in Phb1-SCKO mice. (**a**) Schematics of the rapamycin treatment (left) and mechanism of action of rapamycin (right). (**b**) Representative western blot of sciatic nerve lysates demonstrating that rapamycin treatment reduces the phosphorylation of mTORC1 targets (S6 and 4E-BP1) in Phb1-SCKO mice. (**c**) Quantification of (b). N=5-6 animals per group. Two-way ANOVA corrected for multiple comparisons using the Holm-Sidak method. p-4E-BP1: F (1, 19) interaction = 4.462, p=0.0481; F (1, 19) rapa = 5.509, p=0.0299; F (1, 19) group = 34.16, p<0.0001. 4E-BP1: F (1, 19) interaction = 0.0038, p=0.9514; F (1, 19) rapa = 0.618, p=0.4414; F (1, 19) group = 33.58, p<0.0001. p-S6: F (1, 19) interaction = 7.735, p=0.0119; F (1, 19) rapa = 12.96, p=0.0019; F (1, 19) group = 7.101, p=0.0153. S6: F (1, 19) interaction = 0.01436, p=0.9059; F (1, 19) rapa = 1.373, p=0.2558; F (1, 19) group = 0.8633, p=0.3645. (**d**) Representative tibial nerve sections of the four experimental groups. (**e**) Quantitative analysis of morphological parameters. Rapamycin treatment is able to reduce the number of demyelinated axons and SCs degrading myelin (myelinophagy), as well as increase the number of myelinated fibers in nerves of Phb1-SCKO animals. Insets: Representative images of demyelinated axons (arrows), degenerating axon (arrowhead) and myelinophagy (star). N=5-6 animals per group. Two-way ANOVA corrected for multiple comparisons using the Holm-Sidak method. Myelinated: F (1, 19) interaction = 10.60, p=0.0042; F (1, 19) rapa = 14.05, p=0.0014; F (1, 19) group = 35.45, p<0.0001. (**f**) There is no alteration of myelin thickness (measured by g-ratio = axon diameter/ fiber diameter). N=5-6 animals per group. Two-way ANOVA corrected for multiple comparisons using the Holm-Sidak method. F (1, 20) interaction = 0.1384, p=0.7137; F (1, 20) rapa = 0.01424, p=0.9062; F (1, 20) group = 0.01779, p=0.8952. (**g**) Rapamycin is also able to ameliorate the nerve conduction velocity of mice lacking *Phb1*, but has no effect on CMAP amplitude. N=4-6 animals per group. Two-way ANOVA corrected for multiple comparisons using the Holm-Sidak method. NCV: F (1, 16) interaction = 46.01, p<0.0001; F (1, 20) rapa = 0.01424, p=0.9062; F (1, 20) group = 0.01779, p=0.8952. Amplitude: F (1, 16) interaction = 5.966, p=0.0266; F (1, 16) rapa = 0.2774, p=0.6057; F (1, 16) group = 25.98, p=0.0001. (**h**) Phb1-SCKO mice treated with rapamycin show a trend toward improved performance in the rotarod. N=6 animals per group. Two-way ANOVA corrected for multiple comparisons using the Holm-Sidak method. F (9, 60) interaction = 3.038, p = 0.0047; F (3, 60) time = 58.01, p < 0.0001; F (3, 20) group = 24.84, p < 0.0001. * p<0.05, ** p<0.01, *** p<0.001.

Similar to the studies with JUN, we also evaluated if other pathways of interest could be modulated by mTORC1. Treatment with rapamycin results in a trend toward reduction of JUN expression in Phb1-SCKO mice (**Supplementary Figures 7A and 7B**). It also significantly reduces the protein levels of BiP and p-eIF2α (**Supplementary Figures 7A and 7B**), and the mRNA level of *Ddit3* (**Supplementary Figure 7C**). Therefore, mTORC1 may be a central pathway modulating JUN and the ISR in Phb1-SCKO mice.

## 3. Discussion

About a third of all patients with mitochondrial genetic disorders develop peripheral neuropathies (Pareyson et al., 2013). Most commonly, these patients show axonal degeneration, but, when demyelination is present, the alterations in mitochondrial morphology concentrate in SCs rather than axons (Ino and Iino, 2017). Moreover, many recent reports demonstrate the importance of SC mitochondria in myelin homeostasis in the PNS (Viader et al., 2011, Funfschilling et al., 2012, Niemann et al., 2014, Wang et al., 2016, Della-Flora Nunes et al., 2020). Nevertheless, the SC adaptations to mitochondrial dysfunction and the connection with demyelination remain incompletely understood. Viader et al. (2013) showed that ablation of the mitochondrial transcription factor *Tfam* in SCs triggers the ISR and dysregulates lipid metabolism causing a shift towards lipid degradation. The ISR was proposed to be maladaptive in this context, and this was hypothesized to underly the severe peripheral neuropathy in these mice (Viader et al., 2013). However, we recently demonstrated that the ISR is actually a beneficial response in the context of mitochondrial damage in SCs (Della-Flora Nunes et al., 2020). Therefore, the mechanism proposed by Viader et al. is likely incomplete.

Here we report on a new mechanism involving mTORC1 and JUN that could provide a link between mitochondrial dysfunction in SCs and demyelination. Ablation of the mitochondrial protein PHB1 in SCs in mice results in continuous activation of JUN and mTORC1. Moreover, we demonstrate that mTORC1 can be activated *in vitro* by direct inhibition of mitochondrial function with oligomycin or antimycin A, while JUN is associated with mitochondrial loss in SCs *in vivo*. This supports the hypothesis that JUN and mTORC1 are involved in the SC adaptation to mitochondrial damage. The c-Jun N-terminal Kinase (JNK) signaling pathway has been reported to modulate mitochondrial respiration and production of reactive oxygen species (ROS) (Win et al., 2014, Chambers and LoGrasso, 2011), while mTORC1 is a well-known regulator of metabolism and mitochondrial function (Ramanathan and Schreiber, 2009, de la Cruz López et al., 2019). Therefore, it is possible that mTORC1 and JUN also participate in the response to mitochondrial damage in other cells types. In fact, JUN was suggested to be involved in the activation of the mtUPR in COS-7 cells (Horibe and Hoogenraad, 2007), while mTORC1 was shown to participate in the response to mitochondrial dysfunction in muscle (Khan et al., 2017) and mouse embryonic fibroblasts (Hardy and Pryde, 2020).

SCs present a remarkable plasticity that endows peripheral nerves with the capacity to recover from a variety of insults. After nerve injury, myelinating and non-myelinating SCs convert to a repair-promoting phenotype, allowing degradation of myelin and cell debris and stimulating axon survival and regrowth. Later, these SCs can also differentiate to form new myelin. This entire process is controlled by the transcriptional regulator JUN, whose upregulation requires activation of mTORC1 (for review, see (Jessen and Mirsky, 2019)). Given this particular biology of SCs, we hypothesized that activation of mTORC1 and JUN in the context of mitochondrial damage could inadvertently induce demyelination in Phb1-SCKO mice. In line with this hypothesis, we found a strong association between demyelination (identified by the presence of myelin ovoids) and overactivation of JUN or mTORC1 in these animals. Moreover, deletion of JUN in SCs reduced the demyelination in Phb1-SCKO mice, but also seemed to exacerbate the developmental defects observed in these animals. On the other hand, treatment of Phb1-SCKO mice with rapamycin resulted in almost complete rescue in nerve morphology, while also providing a significant functional benefit in nerve conduction velocity. Thus, both JUN and mTORC1 seem to participate in the demyelination process of Phb1-SCKO mice.

Interestingly, JUN and mTORC1 have also been implicated in other peripheral neuropathies. mTORC1 overactivation may be involved in the focal hypermyelination observed in Charcot–Marie–Tooth disease types 4B1 and 4B2 (CMT4B1 and CMT4B2) (Sawade et al., 2020), while rapamycin treatment is able to ameliorate myelination defects in a mouse model of CMT1A (Nicks et al., 2014). On the other hand, JUN was found to be upregulated in nerve biopsies from patients affected by a variety of peripheral neuropathies (Hutton et al., 2011). Hence, JUN and mTORC1 may underlie key aspects of nerve pathology. Although outside the scope of the current work, one important facet of the pathogenesis of peripheral neuropathies is the impaired trophic support from SCs to axons. JUN was shown to be required to prevent loss of sensory axons in a mouse model of CMT1A (Hantke et al., 2014), while mTORC1 is crucial to trigger a metabolic shift in SCs that supports axonal integrity in the context of subacute acrylamide intoxication (Babetto et al., 2020). Therefore, mTORC1 and JUN may have opposite effects in the SC functions of myelin maintenance and axonal support, and it would be premature to conceive therapeutics targeting these pathways for peripheral neuropathies.

It is unlikely that JUN and mTORC1 are the only pathways involved in demyelination in response to mitochondrial damage in SCs, but we believe they may form an important hub controlling this process together with the ISR. Our pharmacological and genetic approaches revealed that JUN, mTORC1 and ISR are interconnected, with mTORC1 playing a central role and modulating the other two pathways (**Figure 7**). An interesting hypothesis that we would like to explore in the future is that the global control of translation is a key response in SCs downstream of mitochondrial damage. ISR and mTORC1 (in particular its 4E-BP1 arm) are two of the most important pathways regulating cellular translation rates. Activation of the ISR through phosphorylation of eIF2α inhibits global translation and simultaneously promotes the expression of stress response genes in a response coordinated by ATF4. On the other hand, mTORC1 phosphorylates 4E-BP1 relieving its inhibition of eIF4E and promoting translation (Ryoo and Vasudevan, 2017). Therefore, it is interesting that treatment with rapamycin (an approach that should reduce global translation levels) was able to ameliorate the demyelination phenotype of Phb1-SCKO mice, while treatment with ISRIB (an ISR inhibitor that should increase global translation levels) was detrimental for the demyelinating pathology (Della-Flora Nunes et al., 2020). Moreover, it is intriguing that ISRIB was specifically able to modulate the phosphorylation levels of 4E-BP1 and had little effect on the S6K arm of mTORC1. This hypothesis is particularly compelling since, in a model of CMT1B, aberrant activation of translation has already been shown to underlie demyelination in the context of ISR (D’Antonio et al., 2013).

**Figure 7.**
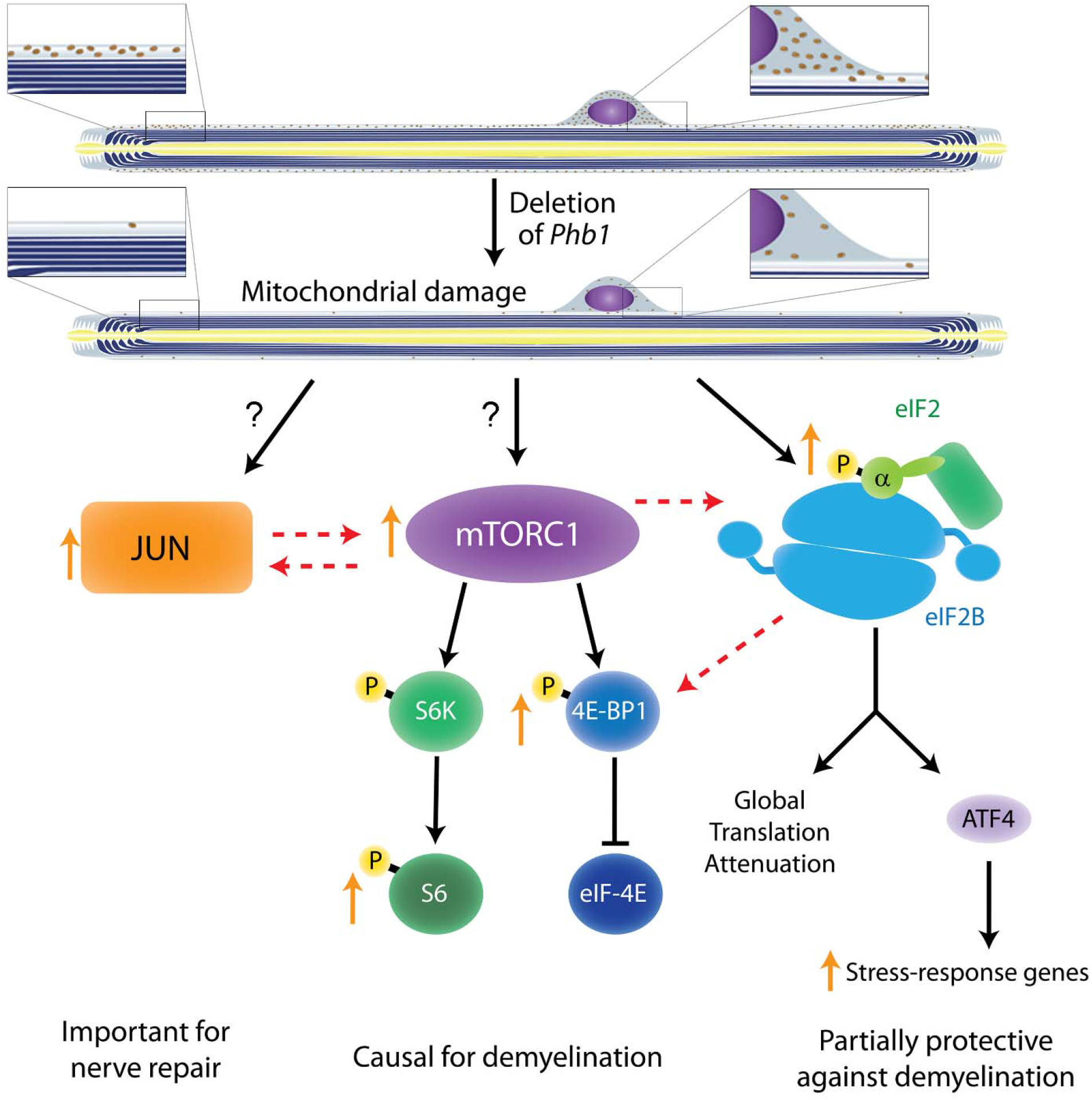
Crosstalk between the pathways investigated in the current study and in (Della-Flora Nunes et al., 2020). Ablation of *Phb1* in SCs leads to severe perturbations to mitochondrial morphology and function, which in turn cause activation of the ISR in myelinating SCs (right). These SCs also upregulate JUN (left) and activate mTORC1 (center), a response that is directly or indirectly associated to the mitochondrial damage. All these pathways partially modulate each other, with mTORC1 playing the most central role and being causal for demyelination. JUN may also participate in demyelination and is key in the nerve repair response, while the ISR is partially protective against demyelination. Discontinuous arrows = partial effects; blunt arrows = inhibition; orange arrows = responses identified in our analyses of Phb1-SCKO mice.

In conclusion, this study reveals a potential new mechanism by which perturbations in SC mitochondria can trigger demyelination. This furthers our understating of how SCs respond to mitochondrial damage and could be relevant in the context of peripheral neuropathies. The link between JUN, mTORC1 and ISR evaluated in our study is also likely to be relevant to other cell types, and may help to advance the general understanding of cellular adaptations elicited in response to mitochondrial dysfunction.

## 4. Materials and methods

### 4.1. Animal models, genotyping and injections

All animal procedures have been approved by the Institutional Animal Care and Use Committee (IACUC) of the Roswell Park Cancer Institute (Buffalo-NY, USA), and followed the guidelines stablished by the NIH’s Guide for the Care and Use of Laboratory Animals and the regulations in place at the University at Buffalo (Buffalo-NY, USA). Animals were housed separated by gender in groups of at most five per cage and kept in a 12h light/dark cycle with water and food *ad libitum*. *Mpz-Cre* and *Phb1* floxed animals were previously described (Feltri et al., 1999, He et al., 2011). Mice were also crossed to the PhAM reporter line (Jackson Laboratories Stock No: 018385) (Pham et al., 2012) and to *Jun* floxed mice (Behrens et al., 2002). Animals were kept in a C57BL/6 and 129 mixed genetic background and analyses were performed from littermates. Animals carrying one or two floxed *Phb1* alleles but no Cre were used as controls, other than for the experiments with PhaM mice, where Control mice were Phb1^wt/wt^; P0-Cre; PhAM, while Phb1-SCKO mice were Phb1^fl/fl^; P0-Cre; PhAM. No animals were excluded from this study.

Genotyping was performed from genomic DNA as previously described for *Mpz-Cre* (Feltri et al., 1999), *Phb1* floxed animals (He et al., 2011), PhaM (Della-Flora Nunes et al., 2020) and *Jun* (Parkinson et al., 2008). Rapamycin (LC Laboratories # R-5000) was prepared as described previously (Beirowski et al., 2017) and administered intraperitoneally at 10 mg/kg daily from P20 to P40. ISRIB (Cayman chemicals # 16258) was prepared as described previously (Chou et al., 2017) and administered intraperitoneally at 2.5 mg/kg daily from P20 to P40.

### 4.2. Morphological assessments

Morphological analyses were performed as described previously (Della-Flora Nunes et al., 2020) Quantification of morphological parameters in electron micrographs used data of ten randomly selected fields at 2,900X magnification. Data were quantified using the cell counter plugin of ImageJ Fiji v1.52p (Rueden et al., 2017, Schindelin et al., 2012).

### 4.3. Behavioral and electrophysiological analyses

These experiments were performed as reported before (Della-Flora Nunes et al., 2020).

### 4.4. Cell culture

Primary rat SCs were prepared as described previously (Brockes et al., 1979) and were not passaged more than four times. Cells were maintained in media containing high glucose DMEM (4.5 g/L glucose) supplemented with 10% fetal bovine serum (FBS), 2mM L-glutamine, 100 U/mL penicillin, 100 μg/mL streptomycin, 2 ng/ml Nrg1 (human NRG1-β1 extracellular domain, R&D Systems), and 2 μM forskolin. For the induction of mitochondrial stress, we prepared stock solutions of 10mM FCCP in ethanol, 40 mM Antimycin A in ethanol and 5 mM Oligomycin in DMSO and stored at −20^°^C until use. For the experiment, 215000 primary rat SCs were plated in each well of a 12-well dish. Two days later, media was exchanged to SC media containing 5 μM FCCP, 2.5 μM oligomycin (Oligo), 10 μM antimycin A (AA) or an equivalent dose of vehicle. For the seven-day treatment, SC media was exchanged every other day with media containing a fresh dilution of the drugs. At the end of the experiment, protein was extracted and western blot was carried out as described below.

### 4.5. Western blot

Western blots were performed as described previously (Della-Flora Nunes et al., 2020). The following primary antibodies were used: Opa1 1:500 (BD Biosciences # 612606), β-tubulin 1:5000 (Novus Biologicals #NB600-936), GAPDH 1:5000 (Sigma #G9545), eIF2α 1:500 (Cell signaling #5324), p-eIF2α 1:500 (Cell signaling #3398), Bip 1:500 (Novus Biologicals #NB300-520), p-4E-BP1 1:500 (Cell signaling #2855), 4E-BP1 1:500 (Cell signaling #9644), p-S6 1:500 (Cell signaling #4858), S6 1:500 (Cell signaling #2217), JUN 1:500 (Cell signaling #9165), p-AKT (Cell signaling #9271); AKT (Cell signaling #9272); MLKL (Abgent #AP14272b); LC3 1:500 (Cell signaling #12741); Atg7 1:500 (Cell signaling #8558); p62 1:500 (Enzo Life Sciences #BML-PW9860). GAPDH or β-tubulin were used as loading controls. All uncropped blots are presented in **Supplementary Figures 8 and 9**.

### 4.6. Immunofluorescence

Immunofluorescence experiments were performed as described (Della-Flora Nunes et al., 2020). For teasing, sciatic or tibial nerves were dissected, fixed in 4% PFA for 30 min, washed with PBS and teased in slides coated with (3-Aminopropyl)triethoxysilane (TESPA; Sigma). Coating with TESPA was achieved by subsequently submerging glass slides in acetone for 1 min, 4% TESPA in acetone for 2 min and two times in acetone for 30 sec each. The teasing procedure consisted of placing a small portion of the nerve in a PBS droplet over the TESPA-coated slide, followed by careful mechanical separation of individual fibers, first using insulin syringes (0.3 ml 31 G x 8 mm) and then using modified insulin syringes containing insect pins (Fine science tools #26002-10) attached to their needle. The following primary antibodies were used: rabbit anti-JUN 1:200 (Cell signaling #9165), rabbit anti-p-S6 1:200 (Cell signaling #4858), chicken anti-P0 1:300 (Aves #PZO0308) and mouse ant-MBP 1:300 (Millipore #MAB384). Images from teased fibers were acquired at 40X magnification and 1.5X zoom using a confocal microscope Leica SP5II running the LAS AF 2.7.9723.3 software (Leica). Quantifications were performed using ImageJ Fiji v1.52p (Rueden et al., 2017, Schindelin et al., 2012). Four to five fields per animal were analyzed.

### 4.7. RNA extraction and qRT-PCR analyses

RNA was isolated and reverse-transcribed as published (Poitelon et al., 2016). qRT-PCR was performed as reported previously (Della-Flora Nunes et al., 2020). Primers for JUN targets were as described previously (Arthur-Farraj et al., 2012).

### 4.8. Statistical analyses

Experiments were not randomized, but data collection and analyses were performed blind to the conditions of the experiments and genotype of the mice. However, due to the severity of the phenotype, in some analyses it was not possible to completely prevent investigators from identifying if the animal was WT or mutant. No data were excluded from the analyses. No power analysis was performed, but our sample sizes are similar to those generally used in the field. The statistical test used in each analysis is reported in the legend of each figure. Data are presented as mean ± s.e.m. P-values < 0.05 were considered to represent a significant difference, while 0.05<p < 0.1 was considered to represent a trend. Data were analyzed using GraphPad Prism 6.01.

## 5. Data availability

The data supporting the findings of this study are available within the article and its supplementary information files. All original data are available from the corresponding author upon reasonable request.

## 7 Acknowledgements

We thank Dr. Erwin F Wagner (Medical University of Vienna) for the *Jun* floxed mice. This work was funded by grant NIH-NINDS-R01NS100464 (to M.L.F.). Generation of the *Phb1* floxed animals was originally supported by National Institutes of Health Grants HD08818 and HD07857 to B.W.O.

## 8. Author contributions

G.D.N. and M.L.F. designed research and interpreted data; G.D.N. carried out the majority of the experiments; E.H. prepared and processed the tissue for morphological analyses; B.H. and B.W.O. provided the *Phb1* floxed mice; G.D.N. and M.L.F. wrote the manuscript; E.R.W., Y.P., L.W., B.W.O. and B.H. analyzed the data and critically reviewed the manuscript.

