## Supplementary Figures for "mTORC1 and JUN are activated after deletion of Prohibitin 1 in Schwann cells and may link mitochondrial dysfunction to demyelination"

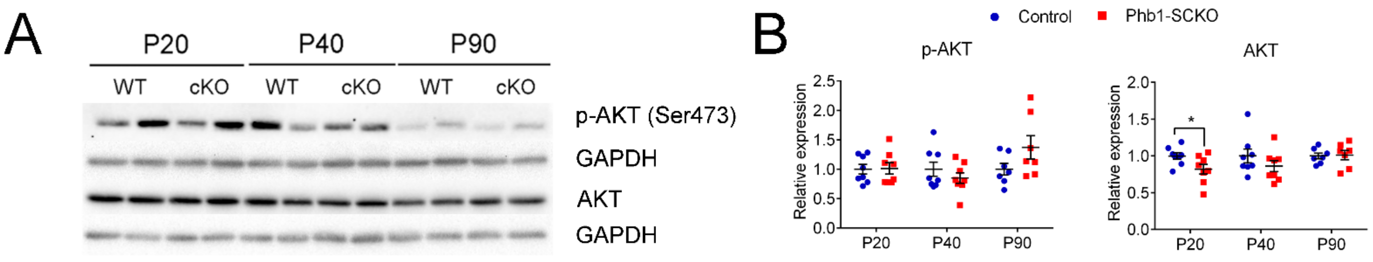


**Supplementary Figure 1**. Akt phosphorylation is not significantly altered by deletion of *Phb1* in SCs. (**a**) Representative western blot from sciatic nerve lysates. (**b**) Quantification of the western blot reveals only minor differences in AKT and p-AKT expression between the genotypes. N=7-8 animals per genotype. Unpaired two-tailed t-test. p-AKT [P20 (t=0.113, df=14), P40 (t=0.999, df=14), P90 (t=1.684, df=12)]; AKT [P20 (t=2.25, df=14), P40 (t=1.62, df=14), P90 (t=0.166, df=12)]. * $p<0.05$

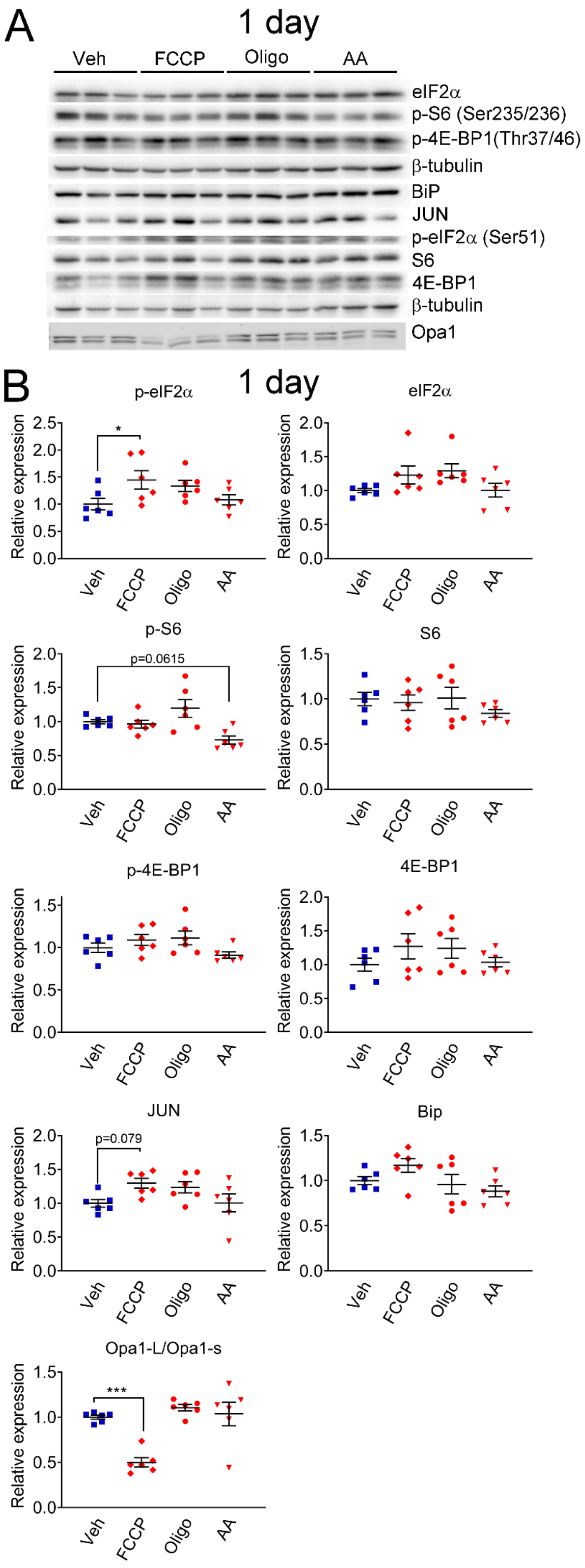


**Supplementary Figure 2**. Short-term mitochondrial perturbation in SCs *in vitro* lead to activation of the ISR. (**a**) Representative western blot from cell lysates of primary rat SCs treated with 5 µM FCCP, 2.5 µM oligomycin (Oligo), 10 µM antimycin A (AA) or vehicle (Veh) for one day. Short-term FCCP induces the integrated stress response (ISR), as seen by elevation of p-eIF2α. (**b**) Quantification of the experiments in (a). N=6 wells per condition. One-way ANOVA corrected for multiple comparisons with the Dunnett method. $F_{(3, 20)}$ p-eIF2α = 1.327, p=0.0568; $F_{(3, 20)}$ eIF2α = 2.355, p=0.1025; $F_{(3, 20)}$ p-S6 = 5.928, p=0.0046; $F_{(3, 20)}$ S6 = 0.842, p=0.487; $F_{(3, 20)}$ p-4E-BP1 = 2.236, p=0.1155; $F_{(3, 20)}$ 4E-BP1 = 1.078, p=0.381; $F_{(3, 20)}$ JUN = 2.911, p=0.0597; $F_{(3, 20)}$ Bip = 2.528, p=0.0864; $F_{(3, 20)}$ Opa1-L/Opa1-s = 14.5, $p<0.0001$. * $p<0.05$, *** $p<0.001$.

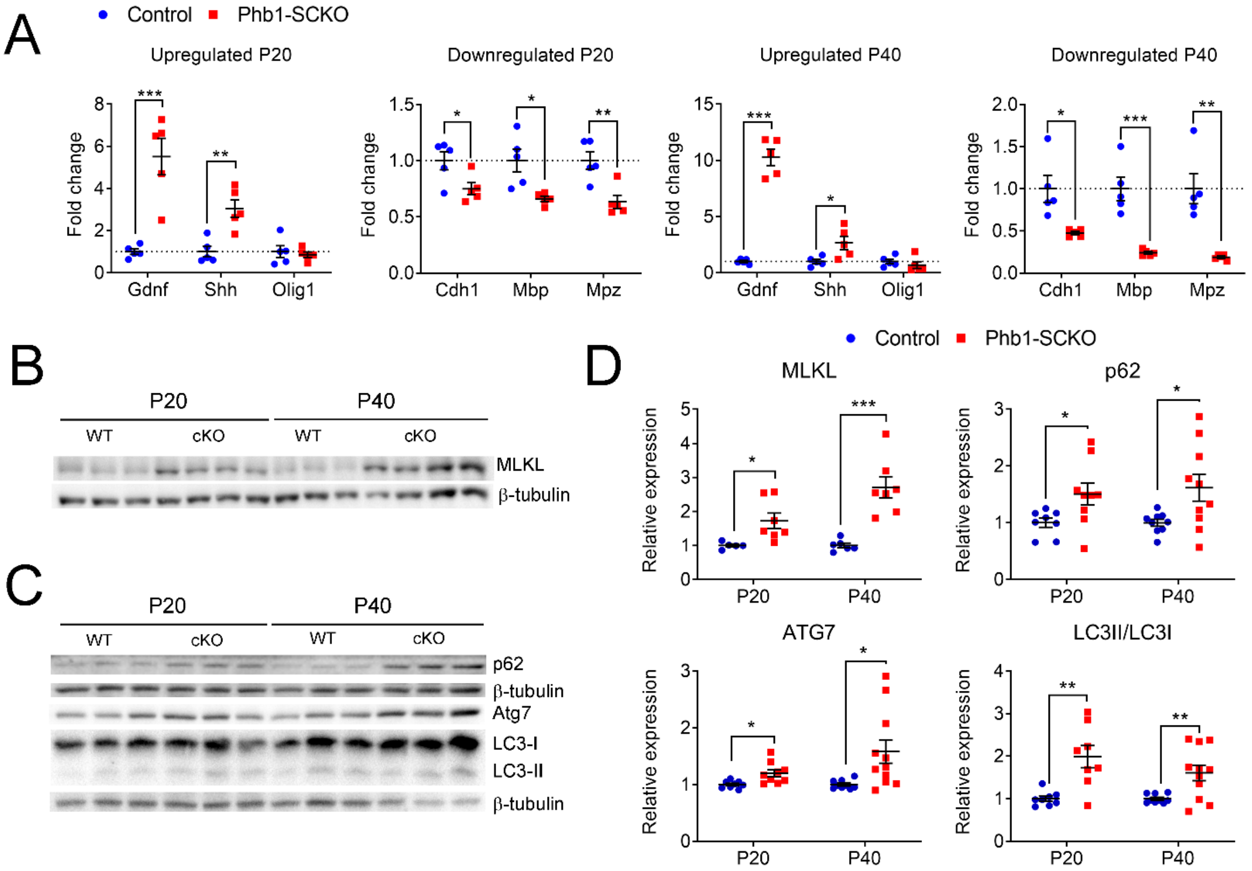


**Supplementary Figure 3.** Phb1-SCKO mice show an overactive myelin breakdown machinery. (**a**) qRT-PCR from sciatic nerves showing that multiple JUN targets have their expression altered in the expected direction when *Phb1* is deleted in SCs. Olig1 was the only JUN target analyzed that was not altered in Phb1-SCKO mice. N=5 animals per genotype. Unpaired two-tailed t-test. Up P20 [*Gdnf* (t=5.143, df=8), *Shh* (t=4.158, df=8), *Olig1* (t=0.46, df=8)]; Down P20 [*Cdh1* (t=2.568, df=8), *Mbp* (t=3.26, df=8), *Mpz* (t=3.757, df=8)]; Up P40 [*Gdnf* (t=12.55, df=8), *Shh* (t=2.673, df=8), *Olig1* (t=0.857, df=8)]; Down P40 [*Cdh1* (t=3.27, df=8), *Mbp* (t=5.447, df=8), *Mpz* (t=4.549, df=8)]. (**b-c**) Representative western blots illustrating the upregulation of MLKL (**b**) and of autophagy machinery (**c**) in sciatic nerves of Phb1-SCKO mice. (**d**) Quantification of the experiments in (a) and (b). N=5-11 animals per genotype. Unpaired two-tailed t-test. MLKL [P20 (t=2.63, df=10), P40 (t=4.937, df=11)]; p62 [P20 (t=2.335, df=15), P40 (t=2.418, df=17)]; ATG7 [P20 (t=2.862, df=15), P40 (t=2.539, df=18)]; LC3II/LC3I [P20 (t=3.734, df=14), P40 (t=2.965, df=18)]. * p<0.05, ** p<0.01, *** p<0.001.

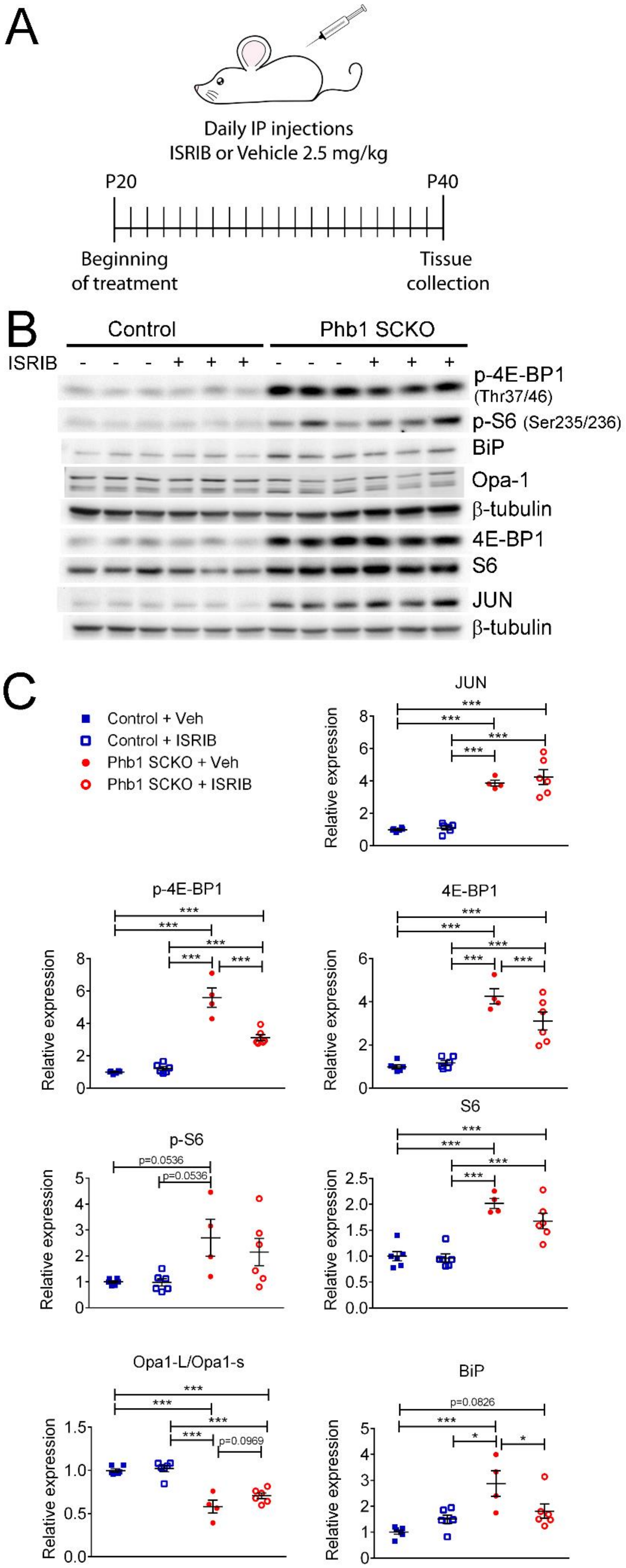


**Supplementary Figure 4.** ISR effect on the other pathways investigated in PHB1-SCKO animals (**a**) Schematic representation of the pharmacological treatment with the ISR inhibitor ISRIB. (**b**) Representative western blots from sciatic nerve lysates demonstrating the effect of ISRIB in P40 Phb1-SCKO mice and controls. (**c**) Quantification of the western blots illustrated in (b). ISRIB only seems to cause small changes in the levels of 4E-BP1 (both total and phosphorylated) and BiP. N=5-6 animals per group. Two-way ANOVA corrected for multiple comparisons using the Holm-Sidak method. JUN: $F_{(1, 18)}$ interaction = 0.267, p=0.611; $F_{(1, 18)}$ ISRIB = 0.823, p=0.376; $F_{(1, 18)}$ group = 125.8, $p<0.0001$. p-4E-BP1: $F_{(1, 18)}$ interaction = 31.17, $p<0.0001$; $F_{(1, 18)}$ ISRIB = 21.56, p=0.0002; $F_{(1, 18)}$ group = 180.5, $p<0.0001$. 4E-BP1: $F_{(1, 18)}$ interaction = 5.778, p=0.027; $F_{(1, 18)}$ ISRIB = 3.133, p=0.0937; $F_{(1, 18)}$ group = 90, $p<0.0001$. p-S6: $F_{(1, 18)}$ interaction = 0.459, p=0.5066; $F_{(1, 18)}$ ISRIB = 0.5113, p=0.484; $F_{(1, 18)}$ group = 13.23, p=0.0019. S6: $F_{(1, 18)}$ interaction = 1.891, p=0.186; $F_{(1, 18)}$ ISRIB = 2.861, p=0.108; $F_{(1, 18)}$ group = 60.64, $p<0.0001$. Opa1-L/Opa1-s: $F_{(1, 18)}$ interaction = 1.719, p=0.206; $F_{(1, 18)}$ ISRIB = 3.334, p=0.0845; $F_{(1, 18)}$ group = 85, $p<0.0001$. BiP: $F_{(1, 18)}$ interaction = 9.354, p=0.0068; $F_{(1, 18)}$ ISRIB = 1.202, p=0.2874; $F_{(1, 18)}$ group = 18.28, p=0.0005. * $p<0.05$, ** $p<0.01$, *** $p<0.001$.

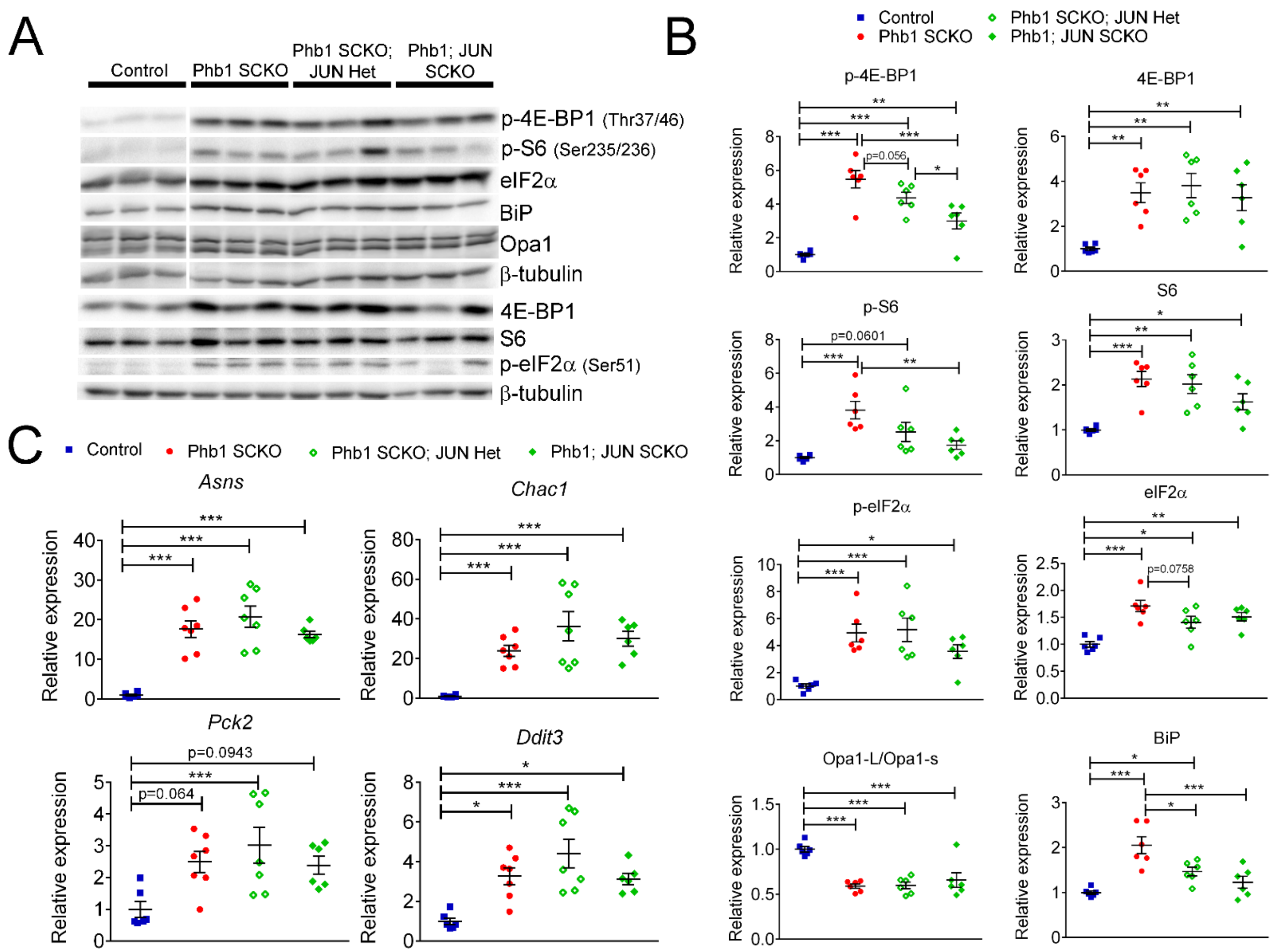


**Supplementary Figure 5.** Effect of JUN in the other pathways altered in PHB1-SCKO animals. (**a**) Representative western blots from sciatic nerve lysates demonstrating the effect of deleting one or two JUN alleles in P40 Phb1-SCKO mice. (**b**) Quantification of the western blots illustrated in (a). N=6 animals per group. One-way ANOVA corrected for multiple comparisons using the Holm-Sidak method. F (3, 20) p-4E-BP1 = 24.74, p<0.0001; F (3, 20) 4E-BP1 = 7.933, p=0.0011; F (3, 20) p-S6 = 8.786, p=0.0006; F (3, 20) S6 = 10.15, p=0.0003; F (3, 20) p-eIF2α = 10.02, p=0.0003; F (3, 20) eIF2α = 11.33, p=0.0001; F (3, 20) Opa1-L/Opa1-s = 15.66, p<0.0001; F (3, 20) BiP = 12.87, p<0.0001. (**c**) qRT-PCR of downstream targets of ATF4 in the ISR in P40 sciatic nerves. Ablation of JUN does not seem to have any effect on the ISR induced by deletion of *Phb1*. N=6-7 animals per group. One-way ANOVA corrected for multiple comparisons using the Holm-Sidak method. F (3, 22) *Asns* = 20.97, p<0.0001; F (3, 22) *Chac1* = 10.76, p=0.0001; F (3, 22) *Pck2* = 4.725, p=0.0108; F (3, 22) *Ddit3* = 8.676, p=0.0005. * p<0.05, ** p<0.01, *** p<0.001.

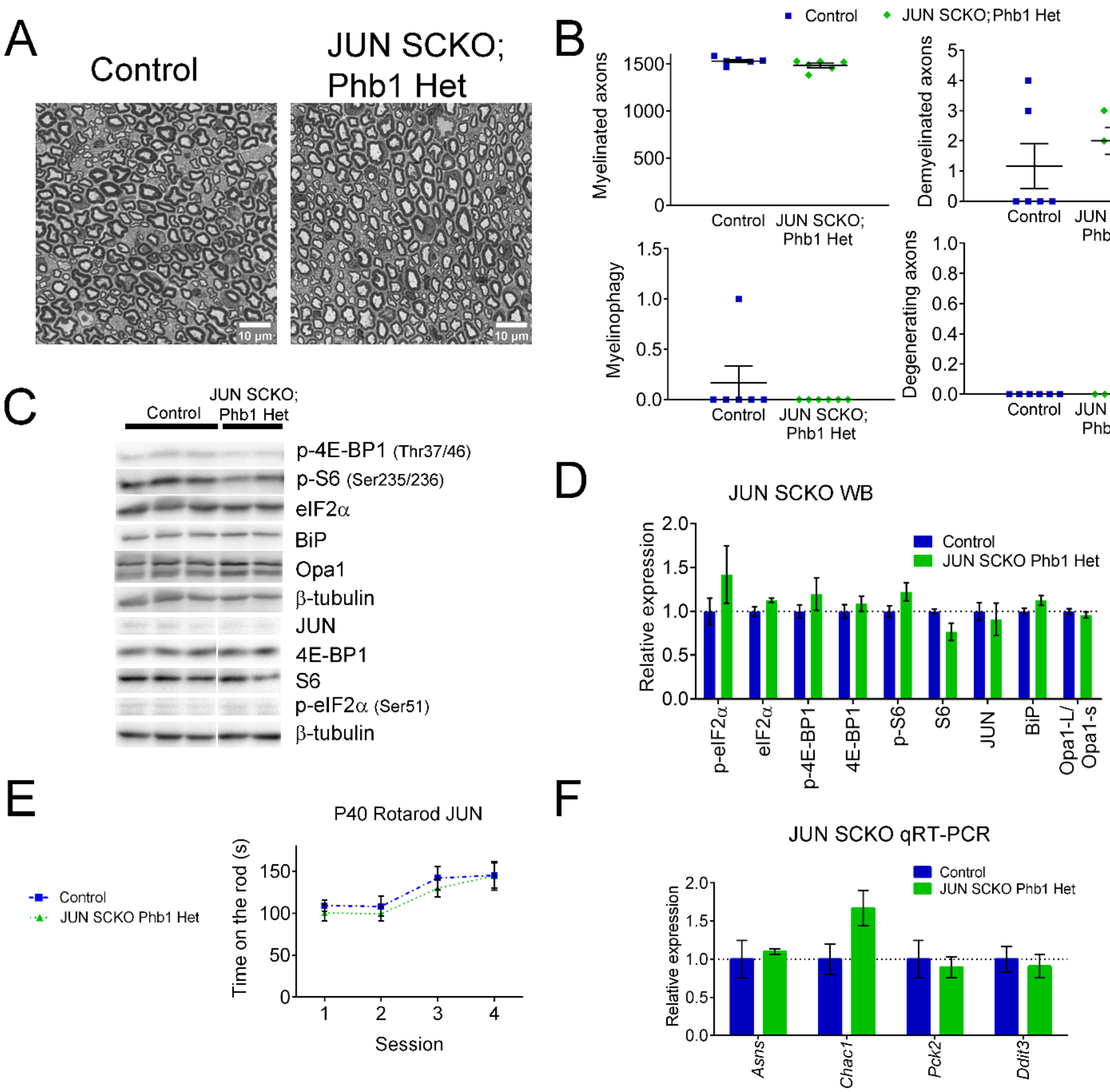


**Supplementary Figure 6.** Deletion of JUN alone has no effect on the parameters analyzed. (**a**) representative semithin images of tibial nerves of control and JUN SCKO; Phb1 Het at P40 (**b**) Morphological parameters quantified from semithin images shows no difference between the two genotypes. N=6 animals per genotype. Unpaired two-tailed t-test. Myelinated (t=1.693, df=10); Demyelinated (t=0.9552, df=10); Degenerating (no t-test possible because all values are identical) (**c**) Representative western blots of sciatic nerves at P40. (**d**) Quantification of (c). N=4-6 animals per group. Unpaired two-tailed t-test corrected for multiple comparisons using the Holm-Sidak method. p-eIF2α (t= 1.3132, df=8); eIF2α (t= 1.895, df=8); p-4E-BP1 (t= 1.134, df=8); 4E-BP1 (t= 0.75, df=8); p-S6 (t= 1.973, df=8); S6 (t= 2.743, df=8); JUN (t= 0.469, df=8); BiP (t= 1.952, df=8); Opa1-L/Opa1-s (t= 0.875, df=8). There are no statistically significant differences between the two groups. (**e**) There is no difference between genotypes in the motor performance evaluated with rotarod. N=6 animals per group. Two-way ANOVA corrected for multiple comparisons using the Holm-Sidak method. F (3, 30) interaction = 0.2048, p = 0.892; F (3, 30) time = 15.31, p < 0.0001; F (1, 10) group = 0.2872, p =0.604. (**f**) qRT-PCR of sciatic nerves at the same age shows no differences between the two groups. N=4-6 animals per group. Unpaired two-tailed t-test corrected for multiple comparisons using the Holm-Sidak method. *Asns* (t= 0.404, df=10); *Chac1* (t= 2.215, df=10); *Pck2* (t= 0.37, df=10); *Ddit3* (t= 0.407, df=10).

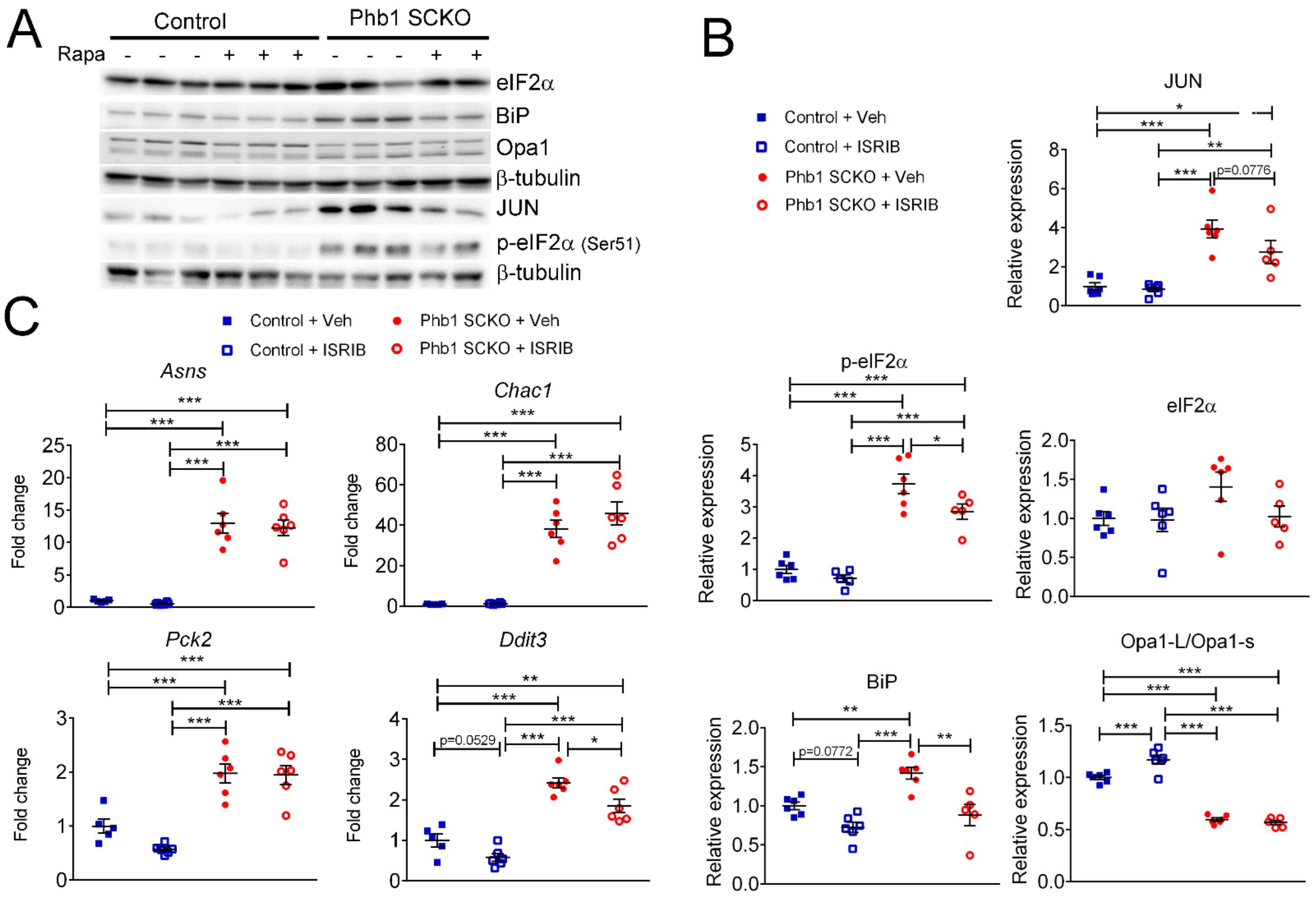


**Supplementary Figure 7**. The role of mTORC1 in the modulation of the other pathways investigated in this work. (**a**) Representative western blots from sciatic nerve lysates of P40 Phb1-SCKO mice and controls treated with rapamycin or vehicle. (**b**) Quantification of the western blots illustrated in (a). N=5-6 animals per group. Two-way ANOVA corrected for multiple comparisons using the Holm-Sidak method. JUN: F (1, 19) interaction = 1.930, p=0.181; F (1, 19) rapa = 3.276, p=0.0862; F (1, 19) group = 43.56, p<0.0001. p-eIF2α: F (1, 19) interaction = 1.778, p=0.199; F (1, 19) rapa = 6.878, p=0.0173; F (1, 19) group = 117.7, p<0.0001. eIF2α: F (1, 19) interaction = 1.538, p=0.23; F (1, 19) rapa = 0.1899, p=0.0173; F (1, 19) group = 2.339, p=0.143. BiP: F (1, 19) interaction = 2.442, p=0.135; F (1, 19) rapa = 23.85, p=0.0001; F (1, 19) group = 11.97, p=0.0026. Opa1-L/Opa1-s: F (1, 19) interaction = 13.16, p=0.0018; F (1, 19) rapa = 6.821, p=0.0169; F (1, 19) group = 339.7, p<0.0001. (**c**) qRT-PCR of downstream targets of ATF4 in the ISR in P40 sciatic nerves. There is no effect of rapamycin treatment, with the exception of an inhibition of the expression of *Ddit3*. N=5-6 animals per group. Two-way ANOVA corrected for multiple comparisons using the Holm-Sidak method. *Asns*: F (1, 19) interaction = 0.0219, p=0.884; F (1, 19) rapa = 0.31, p=0.5842; F (1, 19) group = 132.4, p<0.0001. *Chac1*: F (1, 19) interaction = 0.9956, p=0.331; F (1, 19) rapa = 1.157, p=0.296; F (1, 19) group = 119.7, p<0.0001. *Pck2*: F (1, 19) interaction = 1.929, p=0.181; F (1, 19) rapa = 2.598, p=0.123; F (1, 19) group = 67.51, p<0.0001. *Ddit3*: F (1, 19) interaction = 0.2831, p=0.601; F (1, 19) rapa = 12.4, p=0.0023; F (1, 19) group = 92.39, p<0.0001. * p<0.05, ** p<0.01, *** p<0.001.

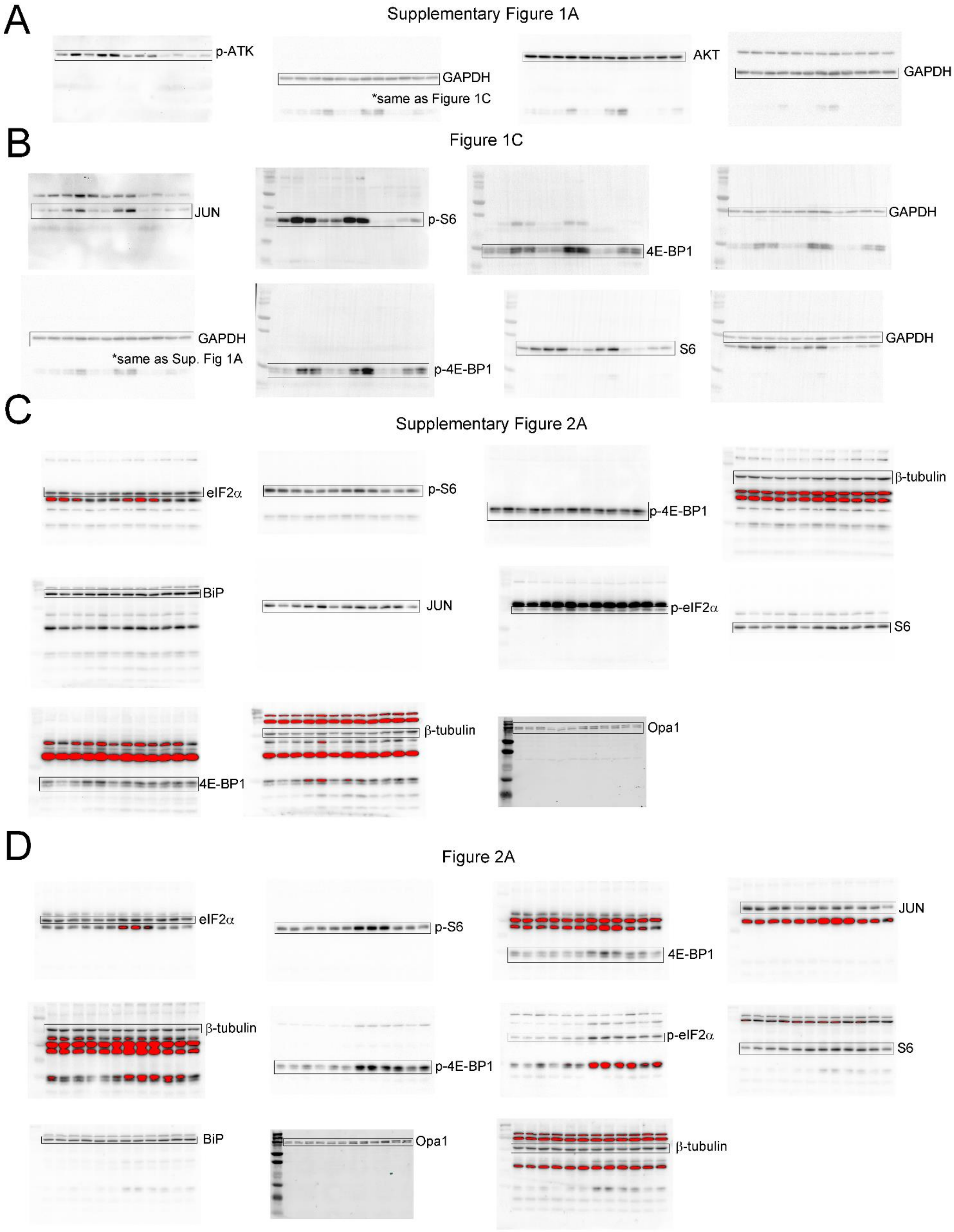


**Supplementary Figure 8.** Part 1 of uncropped blots for representative images of western blots performed in this study.

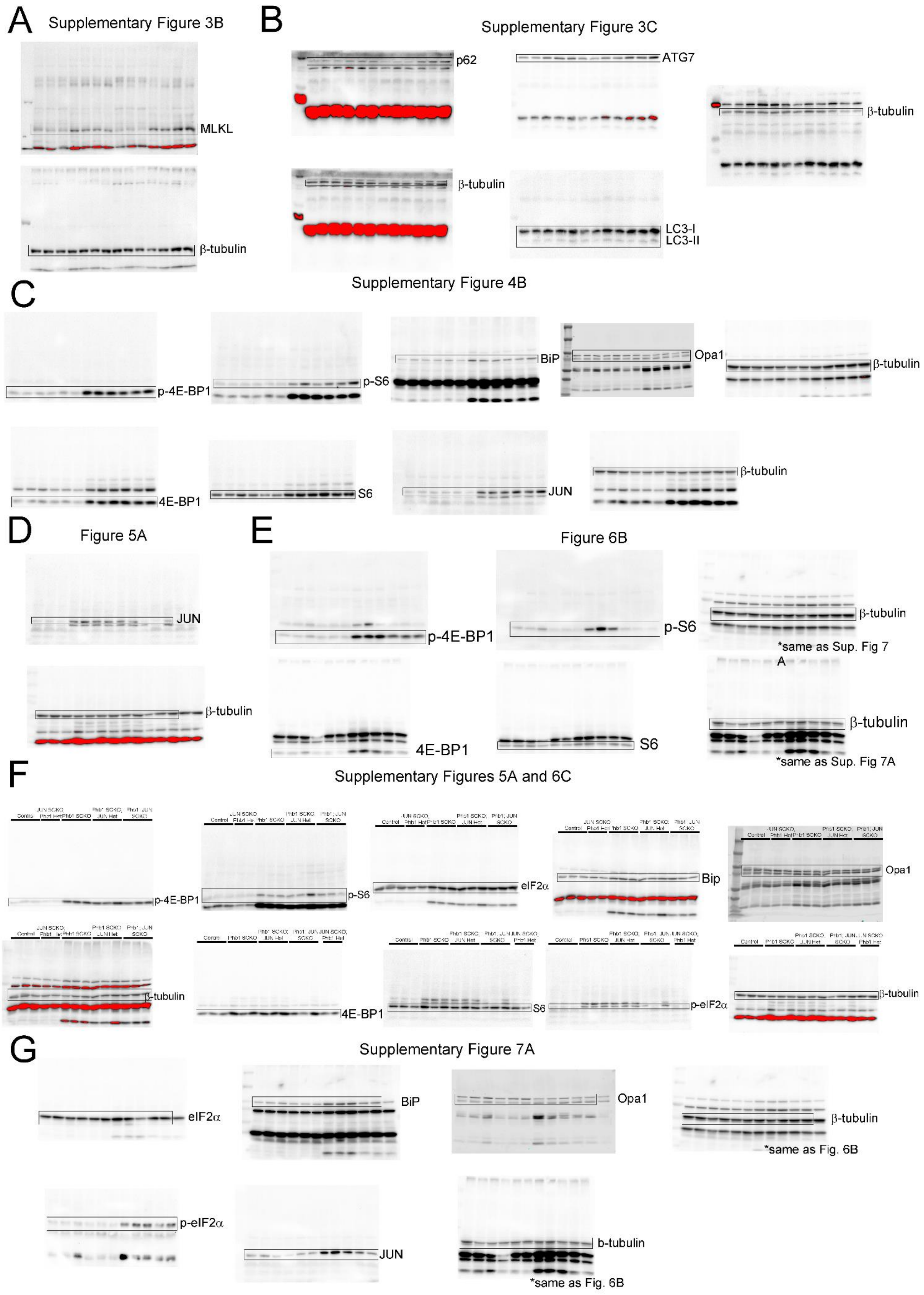


**Supplementary Figure 9.** Part 2 of uncropped blots for representative images of western blots performed in this study.
